# Protein disulfide isomerase disassembles stress granules and blocks cytoplasmic aggregation of TDP-43 in ALS

**DOI:** 10.1101/2024.03.16.585334

**Authors:** Jia-Qi Liu, Hao Liu, Yuying Li, Xiangyi Liu, Li-Qiang Wang, Kan Wang, Zhaofei Yang, Qi Fu, Xiaojiao Xu, Jie Chen, Yingshuang Zhang, Jun Zhou, Weidong Lei, Mengchao Cui, Yi Liang

**Affiliations:** Hubei Key Laboratory of Cell Homeostasis, College of Life Sciences, TaiKang Center for Life and Medical Sciences, Wuhan University, Wuhan 430072, China; Key Laboratory of Radiopharmaceuticals, Ministry of Education, College of Chemistry, Beijing Normal University, Beijing, 100875, China; Department of Neurology, Peking University Third Hospital, Beijing 100191, China; Liaoning Provincial Key Laboratory for Research on the Pathogenic Mechanisms of Neurological Diseases, the First Affiliated Hospital, Dalian Medical University, Dalian 116021, China; Hubei Key Laboratory of Bioinorganic Chemistry & Materia Medica, Hubei Engineering Research Center for Biomaterials and Medical Protective Materials, School of Chemistry and Chemical Engineering, Huazhong University of Science and Technology, Wuhan 430074, China; Institute of Neurology, Sichuan Academy of Medical Sciences-Sichuan Provincial Hospital, Medical School of University of Electronic Science and Technology of China, Chengdu 610072, China; Wuhan University Shenzhen Research Institute, Shenzhen 518057, China; Center for Advanced Materials Research, Beijing Normal University, Zhuhai, 519087, China

## Abstract

Cytoplasmic aggregation of the transactive response DNA-binding protein-43 (TDP-43) in neurons, a pathological feature common to amyotrophic lateral sclerosis (ALS) and frontotemporal lobar degeneration, has been found in some Alzheimer’s patients. Protein disulfide isomerase (PDI) functions as both an enzyme and a molecular chaperone to correct or eliminate misfolded proteins under pathological conditions. Here, we report that TDP-43 is mislocalized to the cytoplasm and colocalizes with PDI in the brain and spinal cord of two ALS patients and the brain of six Alzheimer’s patients compared to controls. TDP-43 selectively recruits wild-type PDI into its phase-separated condensate, which in turn slows down in vitro liquid–liquid phase separation of TDP-43, shifting the equilibrium phase boundary to higher protein concentrations. Importantly, wild-type PDI decreases oxidative stress-induced interaction between TDP-43 and G3BP1 to disassemble stress granules containing TDP-43 in neuronal cells. Wild-type PDI blocks the oxidative stress-induced mislocalization of TDP-43 to the cytoplasm, and blocks subsequent pathological phosphorylation and aggregation of TDP-43. We demonstrate that under pathological stress conditions, wild-type PDI disassembles stress granules, blocks cytoplasmic mislocalization and aggregation of TDP-43, and suppresses mitochondrial damage and TDP-43 toxicity. In the presence of abnormal forms of PDI, however, PDI loses its activity, and stress granules containing TDP-43 are assembled into amyloid fibrils, resulting in mitochondrial impairment and neuronal cell death in ALS patients and some Alzheimer’s patients.

**Teaser:** PDI disassembles SGs, blocks cytoplasmic mislocalization and aggregation of TDP-43, and suppress TDP-43 toxicity in ALS.

## INTRODUCTION

The transactive response DNA-binding protein-43 (TDP-43), a highly conserved, ubiquitously expressed nuclear DNA/RNA-binding protein, has multifaceted functions in regulating RNA metabolism, alternative splicing, miRNA processing, and translation of mRNA (*1*). Under pathological stress conditions, TDP-43 is partly mislocalized from the nucleus to the cytosol and is hyperphosphorylated; cytoplasmic mislocalization and aggregation of TDP-43 in neurons is a common pathological feature in amyotrophic lateral sclerosis (ALS) and frontotemporal lobar degeneration (FTLD-TDP) (*1*−*6*). Cytoplasmic accumulation of Cu, Zn-superoxide dismutase (SOD1) in motor neurons is another pathological feature of ALS (*7*−*9*). Cytoplasmic accumulation of TDP-43 is also observed in the brain of some patients with Alzheimer’s disease (AD) (*10*−*18*) or AD transgenic mice (*14*), but the mechanism behind the phenomenon remains unclear. Protein disulfide isomerase (PDI) functions as both an enzyme and a molecular chaperone to catalyze disulfide bond formation, isomerization, and reduction under physiological conditions and to correct or eliminate misfolded proteins under pathological conditions (*19*−*24*). PDI is mainly located at the endoplasmic reticulum (ER) of cells; under ER stress condition, this ER-resident protein is partly mislocalized from the ER to the cytosol, named PDI ER-to-cytosol reflux (*22*, *24*−*29*). PDI family is implicated in several diverse neurodegenerative diseases, ranging from AD to ALS and Parkinson’s disease (*21*−*24*). However, the exact role of PDI in neurodegenerative diseases is unknown.

Proteins such as RNA-binding proteins tend to form supramolecular assemblies called membrane-less organelles like stress granules via liquid–liquid phase separation (LLPS) of proteins in cells to perform key functions (*30*−*33*). The liquid droplets formed by biological macromolecules, called biomolecular condensates, have fusion properties (*31*, *34*−*40*). Because TDP-43 contains a low-complexity domain in its C-terminal domain (*6*, *36*−*39*, *41*−*43*), it undergoes LLPS in vitro and in cells and forms protein condensates (*35*−*39*, *41*−*45*). TDP-43 liquid-phase condensation is modulated by small molecules, molecular chaperones (Hsp70 and HSPB1), pathological phosphorylation, pathological mutations, RNA (mRNA and yeast total RNA), and other factors (*35*−*39*, *41*−*45*). However, it is unclear whether stress granules containing TDP-43 are assembled into amyloid fibrils in cells. It also remains unknown whether TDP-43 selectively recruits PDI into phase-separated condensates, which in turn regulates the LLPS of TDP-43.

Here, we report that TDP-43 accumulates and colocalizes with PDI in the cytoplasm of the brain of ALS patients and AD patients compared to controls. We demonstrate that TDP-43 selectively recruits wild-type PDI into its phase-separated condensate, which in turn slows down in vitro LLPS of TDP-43 and significantly weakens the interaction between TDP-43 and G3BP1 to disassemble stress granules in neuronal cells under pathological stress conditions. Moreover, wild-type PDI blocks cytoplasmic aggregation of TDP-43 and suppresses mitochondrial damage and TDP-43 toxicity. Our findings provide insights into the regulation of PDI on pathological functions of cytoplasmic TDP-43 via LLPS of the protein inhibited by wild-type PDI, but not abnormal forms of PDI, which has important implications in ALS etiology.

## RESULTS

### TDP-43 accumulates and colocalizes with PDI in the cytoplasm of the brain of ALS patients and AD patients

We first took confocal images of paraffin brain sections from one ALS patient, six AD patients, three healthy controls, and two controls with Castleman disease and progressive supranuclear palsy (PSP), respectively, and paraffin spinal cord section from another ALS patient (Fig. 1, A and B, and Table 1). The brain and spinal cord sections were doubly immunostained with the anti-TDP-43 antibody (green) and anti-PDI antibody (red), stained with DAPI (blue), and visualized by confocal microscopy. We found that TDP-43 (green) was mislocalized and accumulated not only in the cytoplasm of the brain and spinal cord of the two ALS patients but also in the cytoplasm of the brain of six AD patients with TDP-43 pathology (Fig. 1A). In sharp contrast, TDP-43 (green) was mainly located in the nucleus, but was not mislocalized in the cytoplasm in the brain of five age-matched controls in most fields of view (Fig. 1B). Importantly, PDI (red) was partly mislocalized from the ER to the cytosol, and the cytoplasmic TDP-43 and PDI (orange puncta in the merged images) were colocalized not only in the brain and spinal cord of the two ALS patients but also in the brain of six AD patients with TDP-43 pathology (Fig. 1A). In contrast with this, the colocalization of TDP-43 and PDI were not observed in the brain of these five controls (Fig. 1B). Thus, cytoplasmic TDP-43 accumulated and colocalized with PDI in the brain of ALS patients and AD patients (Fig. 1A). The brain and spinal cord were then analyzed by H&E staining (Fig. 1, C and D). H&E staining of the brain and spinal cord sections from two ALS patients and six AD patients showed that the nuclei were dissolved with cell structure disappearance in many neurons (Fig. 1C). Cell vacuolar degeneration was noted in the spinal cord of an ALS patient (Case 2) (Fig. 1C). By contrast, both the dissolution of a cell nucleus and cell structure disappearance were not observed in the brain of five age-matched controls, showing normal cell structures with intact nucleus (Fig. 1D).

**Fig. 1.**
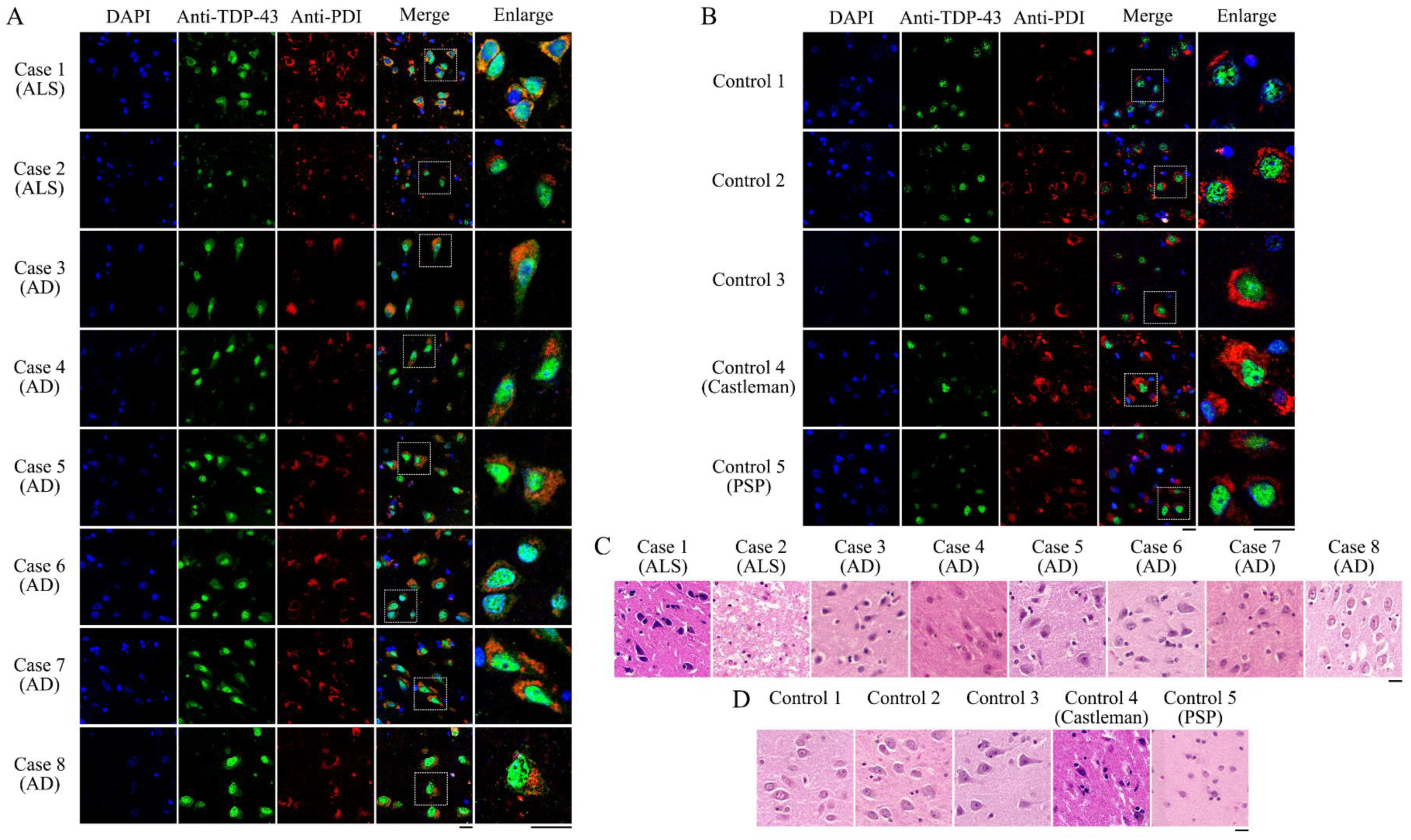
Mislocalization of TDP-43 to the cytoplasm and the colocalization of TDP-43 and PDI were observed in the brain of one ALS patient (Case 1) and six AD patients (Cases 3 to 8) and in the spinal cord of another ALS patient (Case 2), but were not observed in the brain of five age-matched controls (Controls 1 to 5). (**A**) Double immunofluorescence staining of paraffin brain sections from the seven patients (Case 1 and Cases 3 to 8) and paraffin spinal cord section from another ALS patient (Case 2). Shown are nuclei stained with DAPI (blue); signals were detected with antibodies against TDP-43 (green) and PDI (red). The enlarged regions (right) show 9-fold enlarged images from the merged images. Orange puncta indicate colocalization of cytoplasmic TDP-43 and PDI in granules. (**B**) Double immunofluorescence staining of paraffin brain sections from three healthy controls (Controls 1 to 3), one control with Castleman disease (Control 4), and one control with progressive superanuclear palsy (PSP) (Control 5). The experimental conditions are the same as those in (A). (**C**) H&E staining of paraffin brain sections from Case 1 and Cases 3 to 8 and paraffin spinal cord section from Case 2 showed that the nuclei were dissolved with cell structure disappearance in part of neurons. Cell vacuolar degeneration was noted in the spinal cord of Case 2. (**D**) H&E staining of paraffin brain sections from Controls 1 to 5 showed that both the dissolution of a cell nucleus and cell structure disappearance were not observed in the brain of these five controls. Scale bars, 20 μm.

**Table 1.** Clinical details of patients with ALS and Alzheimer’s disease at the time when brain and spinal cord slices were taken.

| Case | Gender | Age | Mean age ± S.D. | TDP-43 pathology | Tau pathology | Amyloid-β pathology | Clinical diagnosis |
| --- | --- | --- | --- | --- | --- | --- | --- |
| ALS and Alzheimer's disease |  |  |  |  |  |  |  |
| 1 | M | 69 | 76.6 ± 11.8 | 4–5 | ND | ND | Sporadic ALS |
| 2 | M | 75 |  | 3–4 | ND | ND | Sporadic ALS |
| 3 | M | 76 |  | ND | V | 5 | Alzheimer's disease |
| 4 | M | 92 |  | ND | V | 4 | Alzheimer's disease |
| 5 | M | 95 |  | ND | III | 1 | Alzheimer's disease |
| 6 | F | 72 |  | ND | I | 1 | Alzheimer's disease |
| 7 | M | 59 |  | ND | III | 4 | Alzheimer's disease |
| 8 | M | 75 |  | ND | V | 6 | Alzheimer's disease |
| Control |  |  |  |  |  |  |  |
| 1 | M | 76 | 74.8 ± 7.3 | ND | 0 | 0 | Healthy individual |
| 2 | F | 70 |  | ND | 0 | 0 | Healthy individual |
| 3 | F | 77 |  | ND | 0 | 0 | Healthy individual |
| 4 | M | 66 |  | ND | 0 | 0 | Castleman disease |
| 5 | M | 85 |  | ND | II | 0 | PSP |

M, male; F, female; ND, not detected.
TDP-43 pathology (NCI/GCI).
ALS, amyotrophic lateral sclerosis; PSP, progressive supranuclear palsy.

### Wild-type PDI slows down LLPS of TDP-43 via specific interactions with TDP-43

We found that TDP-43 accumulated and colocalized with PDI in the cytoplasm of the brain of ALS patients and Alzheimer’s patients (Fig. 1A). Based on these observations, we hypothesized that wild-type PDI specifically interacts with TDP-43 in the cytoplasm of cells under pathological stress conditions. To test this hypothesis, we performed coimmunoprecipitation (co-IP) experiments (Fig. 2A) and took live-cell images of TDP-43-wild-type PDI speck formation (Fig. 2, B to D). Our co-IP experiments visualized by the anti-TDP-43 antibody and anti-HA antibody verified the interaction of TDP-43 with wild-type PDI or dominant-negative PDI (dnPDI), a quadruple cysteine mutant of this enzyme (*21*), in HEK-293T cells transiently co-transfected with TDP-43 and HA-tagged wild-type PDI or HA-tagged dnPDI treated with 1 mM hydrogen peroxide (H_2_O_2_) for 1 h (Fig. 2A). Here, H_2_O_2_ at high concentrations was used to induce oxidative stress and ER stress (*46*−*48*). We then applied a bimolecular fluorescence complementation (BiFC) assay (*49*, *50*) to determine the location of TDP-43-PDI interactions in living cells (Fig. 2D). Schemes of TDP-43-PDI constructs were used in our BiFC assay: TDP-43-EGFP_1-172_ fusion protein (Fig. 2B) and signal peptide-HA-EGFP_155-238_-PDI fusion protein (Fig. 2C). HEK-293T cells transiently expressing both full-length TDP-43-EGFP_1-172_ and EGFP_155-238_-wild-type PDI or EGFP_155-238_-dnPDI constructs were treated with 1 mM H_2_O_2_ for 1 h and cultured for 1 day. Specific interactions between TDP-43 and wild-type PDI or dnPDI in the cytoplasm of living cells under pathological stress (H_2_O_2_-induced oxidative stress) conditions were visualized by our BiFC experiments, and EGFP (green) was observed (Fig. 2D), indicating TDP-43-wild-type PDI speck formation. In sharp contrast, EGFP (green) was not observed in our control experiments (fig. S1, A to D), indicating that no interactions between TDP-43-EGFP_1-172_ and EGFP_155-238_, between EGFP_1-172_ and EGFP_155-238_-wild-type PDI, between EGFP_1-172_ and EGFP_155-238_-dnPDI, or between EGFP_1-172_ and EGFP_155-238_ in living cells were detected by BiFC.

**Fig. 2.**
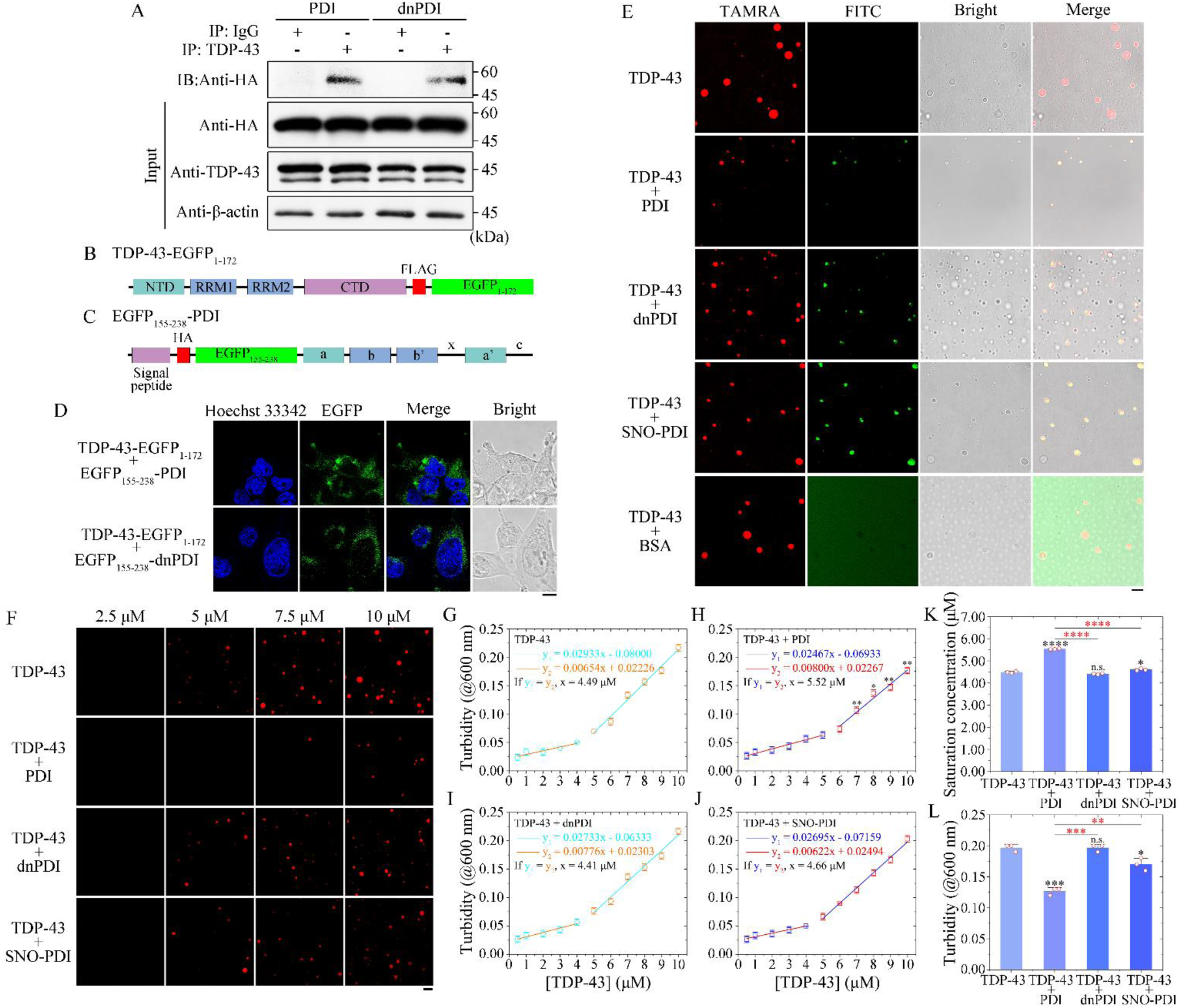
Wild-type PDI slows down LLPS of TDP-43 via specific interactions with TDP-43. (**A**) Co-IP assay to verify the interaction of TDP-43 with PDI in HEK-293T cells transiently co-transfected with TDP-43 and HA-tagged wild-type PDI or HA-tagged dnPDI treated with 1 mM H_2_O_2_ for 1 h. Anti-TDP-43 antibody-binding beads were used for co-IP experiments and then detected by western blotting using anti-HA, anti-TDP-43, and anti-β-actin antibodies. (**B** and **C**) Schemes of TDP-43-PDI constructs used in BiFC assay: TDP-43-EGFP_1-172_ fusion protein (B) and signal peptide-HA-EGFP_155-238_-a-b-b’-x-a’-c fusion protein (C). (**D**) Specific interaction between TDP-43 and PDI in living cells was detected by BiFC. HEK-293T cells transiently expressing both TDP-43-EGFP_1-172_ and EGFP_155-238_-wild-type PDI or EGFP_155-238_-dnPDI constructs were treated with 1 mM H_2_O_2_ for 1 h and cultured for 1 day. Shown are nuclei stained with Hoechst 33342 (blue). EGFP (green) was observed in (D). (**E**) TDP-43 selectively recruits PDI into its phase-separated condensate. Fluorescence images of in vitro phase-separated droplets (red; Merge: yellow) of 10 μM TAMRA-labeled bacterial-purified TDP-43 incubated with Tris-HCl buffer (pH 8.0) containing 10 μM FITC-labeled bacterial-purified wild-type PDI, dnPDI, or SNO-PDI (green) and 10% (w/v) PEG 3350 on ice, and FITC-labeled BSA as a control. (**F**) Regulation of TDP-43 LLPS by PDI. Fluorescence images of 2.5, 5, 7.5, or 10 μM recombinant TDP-43 labeled by TAMRA (red) and incubated with Tris-HCl buffer containing wild-type PDI, dnPDI, or SNO-PDI at 10 μM and 10% PEG 3350 on ice. Scale bars, 10 μm. (**G** to **J**) The dependence of turbidity changes for LLPS of TDP-43 (G), TDP-43 + 10 μM wild-type PDI (H), TDP-43 + 10 μM dnPDI (I), or TDP-43 + 10 μM SNO-PDI (J) on the concentration of TDP-43 ([TDP-43]) was expressed as mean ± SD (with error bars) of values obtained in three independent experiments. The turbidity of TDP-43 condensates was measured at 600 nm at 25 °C. Representative calculation based on turbidity measurements to determine saturation concentrations of TDP-43 or TDP-43 + dnPDI (open circle) and TDP-43 + wild-type PDI or TDP-43 + SNO-PDI (open square). The orange and the red lines are drawn through data points indicating the absence of LLPS, whereas the cyan and the blue lines are drawn through data points in which robust LLPS of TDP-43 occurs. The concentration of protein at which these two lines intersect is an estimation of the saturation concentration. (**K** and **L**) Saturation concentration of TDP-43 (K) and turbidity of TDP-43 condensates (L) (open red circles shown in scatter plots) were expressed as the mean ± SD (with error bars) of values obtained in three biologically independent experiments. TDP-43 + wild-type PDI, *P* = 0.000002 (K) and 0.00012 (L); TDP-43 + dnPDI, *P* = 0.0545 or 0.00000068 (K) and 1.0 or 0.00012 (L); and TDP-43 + SNO-PDI, *P* = 0.0299 or 0.0000000092 (K) and 0.0161 or 0.0029 (L). Statistical analyses were performed using two-sided Student’s *t*-test. Values of *P* < 0.05 indicate statistically significant differences. The following notation is used throughout: \**P* < 0.05; \*\**P* < 0.01; \*\*\**P* < 0.001; and \*\*\*\**P* < 0.0001 relative to control (the saturation concentration for TDP-43 or TDP-43 + wild-type PDI). n.s., no significance.

Given that wild-type PDI specifically interacts with TDP-43 in the cytoplasm of cells under pathological stress conditions (Fig. 2, A and D), we predicted that wild-type PDI might regulate the LLPS of TDP-43 via interaction with the protein. We next used confocal microscopy and turbidity experiments as well as fluorescence recovery after photobleaching (FRAP) (*33*−*40*, *42*−*44*) to test this hypothesis. The freshly bacterial-purified full-length human TDP-43 (residues 1−414), labeled by 5(6)-carboxy-tetramethylrhodamine *N*-succinimidyl ester (TAMRA, red fluorescence) and incubated with Tris-HCl buffer containing 10% PEG 3350 on ice, underwent LLPS in vitro and formed protein condensates (Fig. 2, E and F, and fig. S2, A to C). During the first 20 min after droplet initiation, two small liquid condensates of TDP-43 fused into one larger liquid droplet rapidly within 2 seconds (fig. S2, A and B). After 60 min incubation, however, two small liquid condensates of TDP-43 fused into one larger gel-like/solid-like droplet slowly within 60 seconds (fig. S2C), suggesting that liquid-to-solid phase transitions of TDP-43 occur under such conditions. TDP-43 formed abundant liquid droplets within 20 min after droplet initiation, and protein condensates formed by TDP-43 became much smaller in the presence of low concentrations of wild-type human PDI than in the absence of PDI (Fig. 2, E and F). We took fluorescence images of *in vitro* phase-separated TDP-43 droplets with recombinant, wild-type human PDI, dnPDI, and S-nitrosylated PDI (SNO-PDI) (Fig. 2E). Here, SNO-PDI was assessed by the release of NO, causing the conversion of DAN into the fluorescent compound NAT (fig. S3A). 10 μM TDP-43, labeled by TAMRA (red fluorescence) and incubated with Tris-HCl buffer containing 10 μM FITC-labeled wild-type PDI, dnPDI, or SNO-PDI (green fluorescence) and 10% PEG 3350 on ice, also underwent LLPS in vitro (Fig. 2E). TDP-43 demixed droplets (red; Merge: yellow) coacervated with wild-type PDI, dnPDI, or SNO-PDI (green) were observed by confocal microscopy, with excitation at 561 nm and 488 nm, respectively (Fig. 2E). Importantly, TDP-43 selectively recruited wild-type PDI and dnPDI (or SNO-PDI) but did not recruit nonspecific bovine serum albumin (BSA), a negative control (Fig. 2E), into its phase-separated condensate. Our control experiments demonstrated that wild-type PDI, dnPDI, SNO-PDI, or BSA alone did not form liquid droplets (fig. S3B). Low concentrations of wild-type PDI strongly slowed down in vitro LLPS of TDP-43 (Fig. 2, E and F). In sharp contrast, low concentrations of dnPDI and SNO-PDI, two negative controls and abnormal forms of PDI, did not slow in vitro LLPS of TDP-43 (Fig. 2, E and F). Therefore, TDP-43 selectively recruits wild-type PDI into its phase-separated condensate, which in turn slows down in vitro LLPS of TDP-43. Together, the data showed that wild-type PDI recruited by TDP-43 forms a negative feed-forward loop with TDP-43 to inhibit *in vitro* LLPS of TDP-43.

A generally accepted measure of protein propensity for LLPS is the saturation concentration (*31*, *33*, *34*, *36*, *39*, *44*, *51*−*54*). To assess the effects of PDI on TDP-43 propensity for LLPS, we determined saturation concentrations of TDP-43 in the absence and presence of PDI by measuring the turbidity of TDP-43 condensates at 600 nm as a function of the concentration of TDP-43 (Fig. 2, G−J). The turbidity of a solution does not increase when no LLPS occurs, based on the absorbance at 600 nm (Fig. 2, G−J). We then compared the saturation concentrations of TDP-43 and turbidity of TDP-43 condensates in the absence and presence of PDI (Fig. 2, K and L). Importantly, we found that the saturation concentration of TDP-43 in the presence of wild-type PDI (5.54 ± 0.02 μM, *p* = 0.000002) was significantly higher than that of TDP-43 in the absence of PDI (4.49 ± 0.04 μM), but dnPDI and SNO-PDI did not significantly change the saturation concentration of TDP-43 (4.41 ± 0.03 μM, *P* = 0.0545; 4.60 ± 0.05 μM, *P* = 0.0299) (Fig. 2K). We also found that the turbidity of TDP-43 condensates in the presence of wild-type PDI (0.127 ± 0.006, *p* = 0.00012) was significantly lower than that of TDP-43 condensates in the absence of PDI (0.197 ± 0.006), but dnPDI and SNO-PDI did not significantly change the turbidity of TDP-43 condensates (0.197 ± 0.006, *P* = 1.0; 0.17 ± 0.01, *P* = 0.016) (Fig. 2L). Together, the data showed that wild-type PDI strongly slows down the LLPS of TDP-43 via specific interactions with TDP-43, shifting the equilibrium phase boundary to higher protein concentrations.

We then investigated and evaluated the dynamics of *in vitro* phase-separated droplets of TDP-43 with/without PDI by FRAP (fig. S4, A to E). FRAP of phase-separated TDP-43 droplets without PDI or with two negative controls dnPDI and SNO-PDI revealed fluorescence recovery of (32.6 ± 0.7)% or (31.6 ± 3.6)% and (37.5 ± 4.6)% within 250 s (fig. S4, E and F). In sharp contrast, FRAP of phase-separated TDP-43 droplets coacervated with wild-type PDI revealed much higher fluorescence recovery, (46.7 ± 2.2)%, within 250 s (fig. S4, E and F). According to fig. S4, wild-type PDI significantly enhanced fluorescence recovery. This means that wild-type PDI increases the fluidity of LLPS condensates, possibly through blocking liquid-to-solid transitions in phase-separated TDP-43 condensates. The aforementioned experiments help drive the narrative that wild-type PDI at low concentrations enhances fluorescence recovery and negatively modulates LLPS of TDP-43.

Altogether, these data strongly suggest that the interactions between TDP-43 and wild-type PDI control liquidity and that wild-type PDI enhances TDP-43 mobility via specific interactions with TDP-43. Importantly, wild-type PDI slows down the LLPS of TDP-43 via increasing the saturation concentration for TDP-43 condensation. Therefore, wild-type PDI is a key factor in modulating TDP-43 liquid-phase condensation under pathological stress conditions.

### Wild-type PDI blocks the oxidative stress-induce mislocalization of TDP-43 to the cytoplasm

Increased levels of oxidative stress and ER stress are both observed in the brain of patients with ALS (*55*, *56*). To mimic oxidative stress and ER stress conditions in neuronal cells, we treated both N2a neuroblastoma cells (Fig. 3, A to C) and HEK-293T cells (fig. S5, A to C) with 1 mM H_2_O_2_, and observed a clear shift of endogenous TDP-43 to the cytoplasm of both cell lines (Fig. 3 and fig. S5, A to C). We then compared the amount of endogenous TDP-43 in the cytoplasm and nucleus of N2a cells (Fig. 3, B and C) or HEK-293T cells (fig. S5, B and C) in the absence and presence of transiently expressed PDI. Importantly, we found that under H_2_O_2_-induced oxidative stress conditions, the amount of endogenous TDP-43 in cytoplasm of the two cell lines in the presence of wild-type PDI (0.469 ± 0.012, *p* = 0.000086; 1.19 ± 0.10, *p* = 0.00049) was significantly lower than that in the absence of PDI (0.817 ± 0.035; 1.93 ± 0.07), and the amount of endogenous TDP-43 in nucleus of the two cell lines in the presence of wild-type PDI (1.89 ± 0.14, *p* = 0.00048; 1.54 ± 0.01, *p* = 0.00099) was significantly higher than that in the absence of PDI (0.979 ± 0.069; 1.23 ± 0.06), but dnPDI did not significantly change the amount of endogenous TDP-43 in cytoplasm of the two cell lines (0.794 ± 0.045, *p* = 0.535; 1.95 ± 0.02, *p* = 0.741) and only mildly changed the amount of endogenous TDP-43 in nucleus (1.36 ± 0.13, *p* = 0.0104; 1.30 ± 0.05, *p* = 0.20) (Fig. 3 and fig. S5, B and C). We next took confocal images of N2a cells or HEK-293T cells transiently expressing EGFP-TDP-43 and empty vector pcDNA3.1 treated with 1 mM H_2_O_2_ for 1 h, transiently expressing EGFP-TDP-43 and wild-type PDI treated with 1 mM H_2_O_2_ for 1 h, and transiently expressing EGFP-TDP-43 and dnPDI also treated with 1 mM H_2_O_2_ for 1 h, using N2a cells or HEK-293T cells transiently expressing EGFP-TDP-43 and pcDNA3.1 as a control. The above cells were immunostained with the anti-PDI antibody (red), stained with DAPI (blue), and observed by confocal microscopy (Fig. 3D and fig. S5D). Under the pathological stress condition, EGFP-TDP-43 (green) was partly mislocalized from the nucleus to the cytoplasm of both cell lines (Fig. 3D and fig. S5D). Importantly, we found that under the pathological stress condition, wild-type PDI blocked the mislocalization of EGFP-TDP-43 to the cytoplasm of both cell lines in most fields of view, but dnPDI did not block the mislocalization of EGFP-TDP-43 to the cytoplasm (Fig. 3D and fig. S5D). We once again observed that wild-type PDI specifically interacted with TDP-43 in the cytoplasm of cells under the pathological stress condition (Fig. 3D and fig. S5D). Together, the data showed that wild-type PDI blocked the oxidative stress-induced mislocalization of TDP-43 to the cytoplasm.

**Fig. 3.**
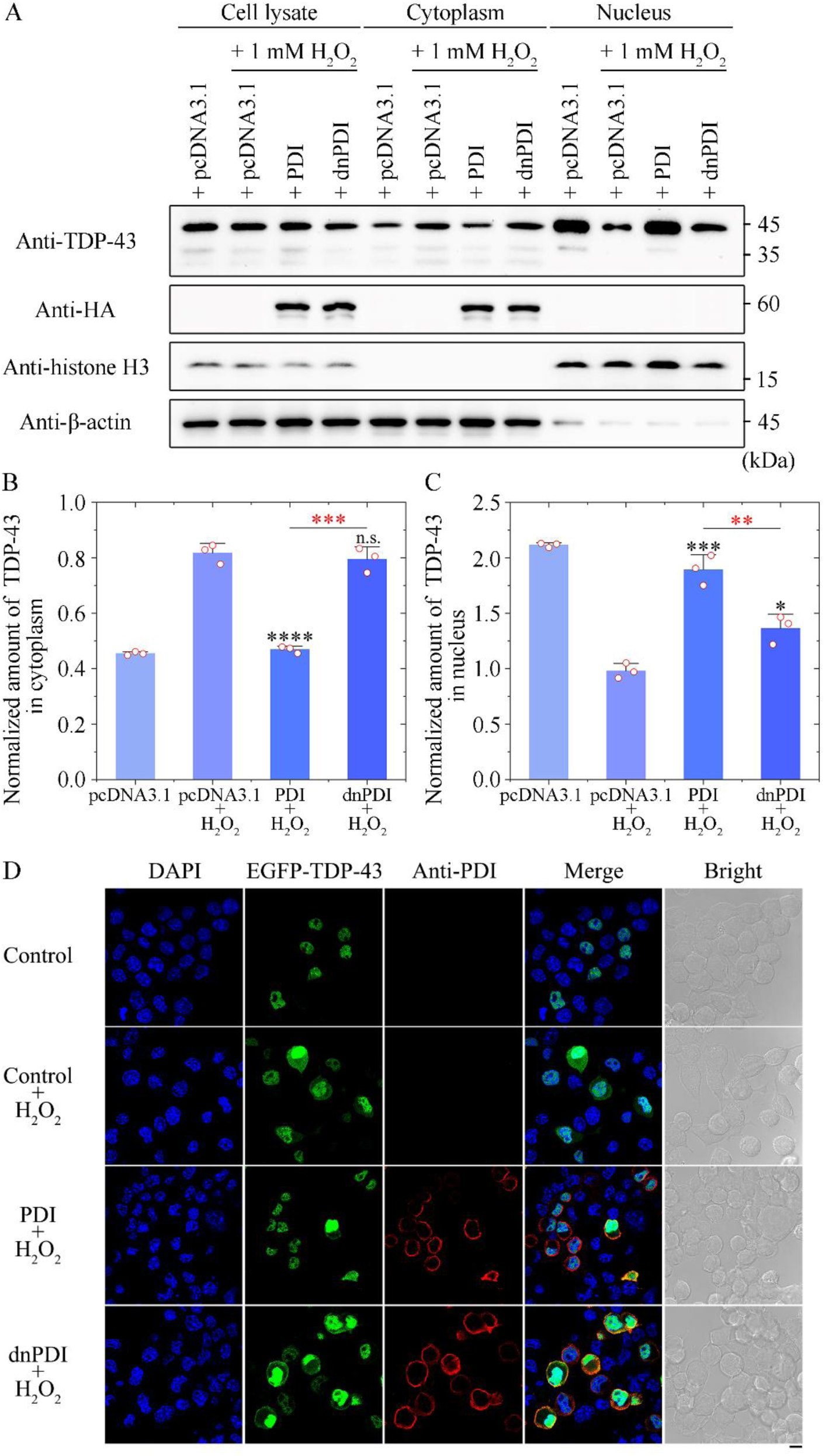
Wild-type PDI blocks the mislocalization of TDP-43 to the cytoplasm induced by H_2_O_2_. (**A**) Western blot for endogenous TDP-43 in the cytoplasmic and nuclear protein fractions and the corresponding cell lysates from N2a cells transiently expressing HA-tagged wild-type PDI or HA-tagged dnPDI treated with 1 mM H_2_O_2_ for 1 h. β-actin and histone H3 served as markers for cytosol and nuclear fractions, respectively, and empty vector pcDNA3.1 as a control. (**B** and **C**) The normalized amount of endogenous TDP-43 in cytoplasm (B) and nucleus (C) of the aforementioned cells (open red circles shown in scatter plots) was expressed as the mean ± SD (with error bars) of values obtained in three independent experiments. Wild-type PDI + H_2_O_2_, *P* = 0.000086 (B) and 0.00048 (C); and dnPDI + H_2_O_2_, *P* = 0.535 or 0.00026 (B) and 0.0104 or 0.0081 (C). Statistical analyses were performed using two-sided Student’s *t*-test. Values of *P* < 0.05 indicate statistically significant differences. The following notation is used throughout: \**P* < 0.05; \*\**P* < 0.01; \*\*\**P* < 0.001; and \*\*\*\**P* < 0.0001 relative to control (pcDNA3.1 + H_2_O_2_ or wild-type PDI + H_2_O_2_). n.s., no significance. (**D**) Immunofluorescence imaging of N2a cells transiently expressing both EGFP-TDP-43 and empty vector pcDNA3.1 (control), transiently expressing both EGFP-TDP-43 and pcDNA3.1 treated with 1 mM H_2_O_2_ for 1 h (control + H_2_O_2_), transiently expressing both EGFP-TDP-43 and wild-type PDI treated with 1 mM H_2_O_2_ for 1 h (wild-type PDI + H_2_O_2_), and transiently expressing both EGFP-TDP-43 and dnPDI also treated with 1 mM H_2_O_2_ for 1 h (dnPDI + H_2_O_2_), using antibody against PDI (red) and staining with DAPI (blue). EGFP-TDP-43 (green) was observed in (D). Scale bar, 10 μm.

### Wild-type PDI decreases oxidative stress-induced interaction between TDP-43 and G3BP1 to disassemble stress granules in neuronal cells

Given that TDP-43 selectively recruits wild-type PDI into its phase-separated condensate, which in turn slows down in vitro LLPS of TDP-43 (Fig. 2), we predicted that wild-type PDI might regulate in vivo LLPS of TDP-43 and thus disassemble stress granules containing TDP-43 in neuronal cells. G3BP1, a molecular switch, triggers phase separation to assemble stress granules, and undergoes K63-linked ubiquitination to modulate stress granule disassembly (*52*, *53*, *57*−*60*). We next used co-IP and confocal microscopy to test this hypothesis. Our co-IP experiments visualized by the anti-G3BP1 antibody, anti-TDP-43 antibody, and anti-HA antibody verified the interaction of endogenous TDP-43 with endogenous G3BP1 in N2a cells transiently expressing empty vector pcDNA3.1, transiently expressing pcDNA3.1 treated with 1 mM H_2_O_2_ for 1 h, transiently expressing wild-type PDI treated with 1 mM H_2_O_2_ for 1 h, and transiently expressing dnPDI also treated with 1 mM H_2_O_2_ for 1 h (Fig. 4A). We then compared the amount of immunoprecipitated G3BP1 in N2a cells in the absence and presence of transiently expressed PDI (Fig. 4B). Importantly, we found that under the H_2_O_2_-induced oxidative stress condition, the amount of immunoprecipitated G3BP1 in N2a cells in the presence of wild-type PDI (0.523 ± 0.041, *p* = 0.00031) was significantly lower than that in the absence of PDI (0.820 ± 0.017), but dnPDI did not significantly change the amount of immunoprecipitated G3BP1 in N2a cells (0.795 ± 0.030, *p* = 0.270) (Fig. 4B). We next took confocal images of N2a cells transiently expressing EGFP-TDP-43 and empty vector pcDNA3.1 treated with 1 mM H_2_O_2_ for 1 h, transiently expressing EGFP-TDP-43 and wild-type PDI treated with 1 mM H_2_O_2_ for 1 h, and transiently expressing EGFP-TDP-43 and dnPDI also treated with 1 mM H_2_O_2_ for 1 h, using N2a cells transiently expressing EGFP-TDP-43 and pcDNA3.1 as a control. The above cells were doubly immunostained with the anti-PDI antibody (red) and anti-G3BP1 antibody (magenta), stained with DAPI (blue), and observed by confocal microscopy (Fig. 4C). Under pathological stress conditions, EGFP-TDP-43 (green) was partly mislocalized from the nucleus to the cytoplasm of N2a cells, and the abundant white puncta observed in confocal images represented the colocalization of TDP-43 and endogenous G3BP1 in stress granules (Fig. 4C). Importantly, we found that under the pathological stress condition, wild-type PDI blocked the mislocalization of EGFP-TDP-43 to the cytoplasm in most fields of view (Fig. 4C) and significantly decreased oxidative stress-induced interaction between TDP-43 and G3BP1 to disassemble stress granules containing TDP-43 in N2a cells (Fig. 4). In sharp contrast, dnPDI did not block the mislocalization of EGFP-TDP-43 to the cytoplasm of N2a cells, and the abundant white puncta observed in confocal images represented the colocalization of TDP-43 and endogenous G3BP1 in the stress granules in the presence of dnPDI (Fig. 4C).

**Fig. 4.**
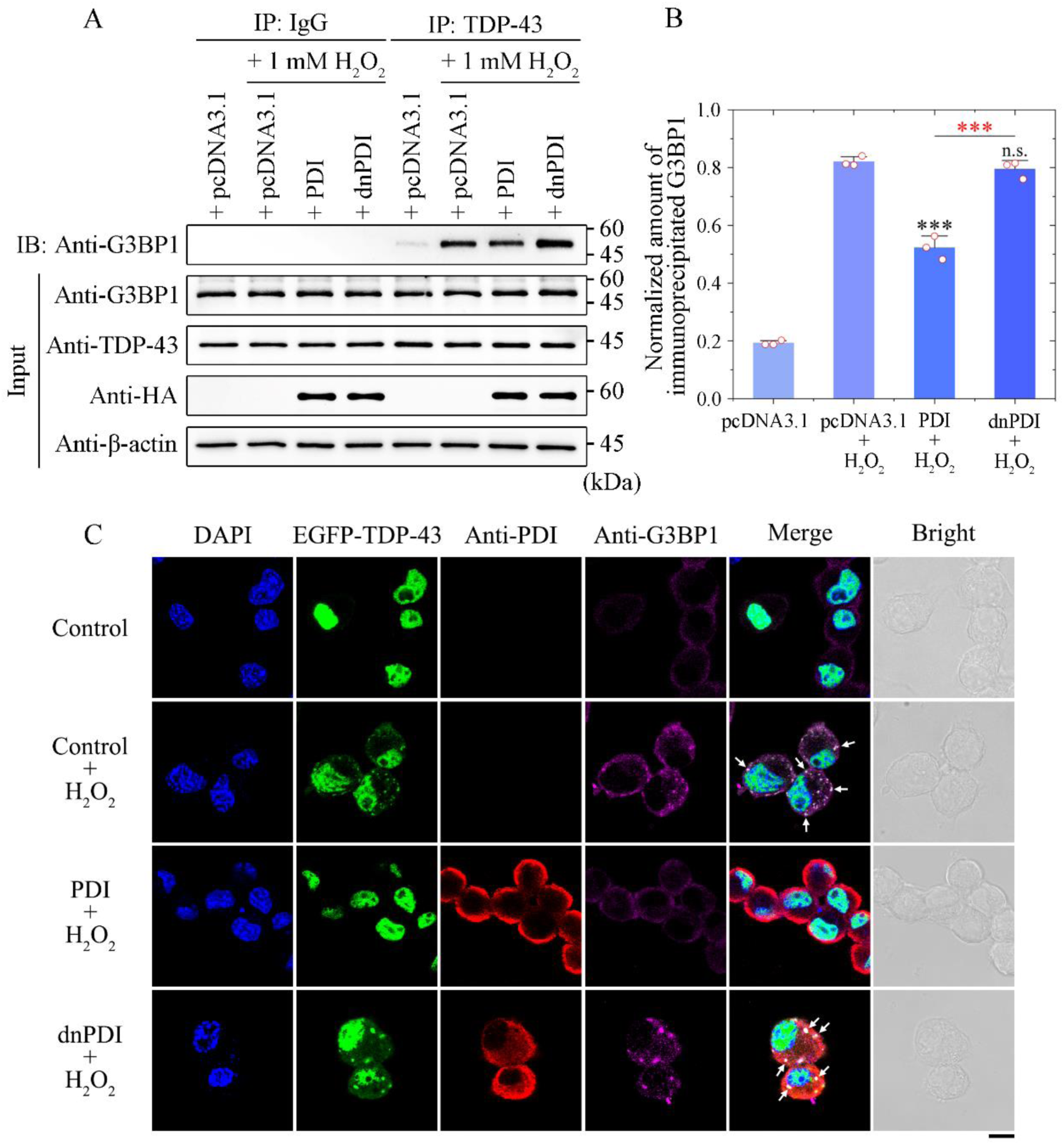
Wild-type PDI decreases H_2_O_2_-induced interaction between TDP-43 and G3BP1 to disassemble stress granules containing TDP-43 in neuronal cells. (**A**) Co-IP assay to verify the interaction of endogenous TDP-43 with endogenous G3BP1 in N2a cells transiently expressing empty vector pcDNA3.1, transiently expressing pcDNA3.1 treated with 1 mM H_2_O_2_ for 1 h, transiently expressing wild-type PDI treated with 1 mM H_2_O_2_ for 1 h, and transiently expressing dnPDI also treated with 1 mM H_2_O_2_ for 1 h. Anti-TDP-43 antibody-binding beads were used for co-IP experiments and then detected by western blotting using anti-G3BP1, anti-TDP-43, anti-HA, and anti-β-actin antibodies. (**B**) The normalized amount of immunoprecipitated G3BP1 in the aforementioned cells (open red circles shown in scatter plots) was expressed as the mean ± SD (with error bars) of values obtained in three independent experiments. Wild-type PDI + H_2_O_2_, *P* = 0.00031; and dnPDI + H_2_O_2_, *P* = 0.270 or 0.00073. Statistical analyses were performed using two-sided Student’s *t*-test. Values of *P* < 0.05 indicate statistically significant differences. The following notation is used throughout: \**P* < 0.05; \*\**P* < 0.01; and \*\*\**P* < 0.001 relative to control (pcDNA3.1 + H_2_O_2_ or wild-type PDI + H_2_O_2_). n.s., no significance. **(C**) Immunofluorescence imaging of N2a cells transiently expressing both EGFP-TDP-43 and pcDNA3.1 (control), transiently expressing both EGFP-TDP-43 and pcDNA3.1 treated with 1 mM H_2_O_2_ for 1 h (control + H_2_O_2_), transiently expressing both EGFP-TDP-43 and wild-type PDI treated with 1 mM H_2_O_2_ for 1 h (wild-type PDI + H_2_O_2_), and transiently expressing both EGFP-TDP-43 and dnPDI treated with 1 mM H_2_O_2_ for 1 h (dnPDI + H_2_O_2_), using antibodies against PDI (red) and G3BP1 (magenta) and staining with DAPI (blue). EGFP-TDP-43 (green) was observed in (C). White arrows indicate colocalization of TDP-43 and endogenous G3BP1 in stress granules. Scale bar, 10 μm.

Altogether, these data demonstrate that TDP-43 selectively recruits wild-type PDI into its phase-separated condensate, which in turn slows down i*n vivo* LLPS of TDP-43 and significantly weakens the interaction between TDP-43 and G3BP1 to disassemble stress granules in neuronal cells under pathological stress conditions.

### Wild-type PDI blocks the oxidative stress-induced pathological phosphorylation and aggregation of TDP-43 to suppress mitochondrial damage and TDP-43 toxicity

Given that TDP-43 selectively recruits wild-type PDI into its phase-separated condensate, which in turn slows down not only *in vitro* LLPS of TDP-43 (Fig. 2) but also *in vivo* LLPS of TDP-43 under pathological stress conditions (Fig. 4), we predicted that wild-type PDI might regulate pathological phosphorylation and aggregation of TDP-43 in neuronal cells under oxidative stress condition. We next employed western blotting, confocal microscopy, and immunogold electron microscopy (*8*, *14*, *28*, *37*, *40*) to test this hypothesis. To test the functional relevance of the observations that we made in vitro and in vivo with TDP-43 LLPS, we used N2a cells or HEK-293T cells transiently expressing empty vector pcDNA3.1 treated with 1 mM H_2_O_2_ for 1 h, transiently expressing wild-type PDI treated with 1 mM H_2_O_2_ for 1 h, and transiently expressing dnPDI also treated with 1 mM H_2_O_2_ for 1 h, using N2a cells or HEK-293T cells transiently expressing pcDNA3.1 as a control. We evaluated pathological phosphorylation of TDP-43 using anti-pSer409/410 antibody, the most common anti-pTDP-43 antibody (*4*, *17*, *18*, *35*, *42*, *43*, *47*). The cell lysates from the above cells were probed by the anti-pTDP-43 antibody, anti-TDP-43 antibody, anti-HA antibody, and anti-β-actin antibody (Fig. 5A and fig. S6A). Treatment of H_2_O_2_ strongly promoted the pathological phosphorylation of endogenous TDP-43 in both cell lines (Fig. 5A and fig. S6A). We then compared the amount of phosphorylated endogenous TDP-43 in the two cell lines in the absence and presence of transiently expressed PDI (Fig. 5B and fig. S6B). Importantly, we found that under the H_2_O_2_-induced oxidative stress condition, the amount of phosphorylated endogenous TDP-43 in both cell lines in the presence of wild-type PDI (0.590 ± 0.045, *p* = 0.00083; 0.608 ± 0.045, *p* = 0.00027) was significantly lower than that in the absence of PDI (1.21 ± 0.11; 0.948 ± 0.018), but dnPDI did not significantly change the amount of phosphorylated endogenous TDP-43 in the two cell lines (1.106 ± 0.058, *p* = 0.211; 0.947 ± 0.050, *p* = 0.977) (Fig. 5B and fig. S6B). We next took confocal images of N2a cells or HEK-293T cells transiently expressing EGFP-TDP-43 and empty vector pcDNA3.1 treated with 1 mM H_2_O_2_ for 1 h, transiently expressing EGFP-TDP-43 and wild-type PDI treated with 1 mM H_2_O_2_ for 1 h, and transiently expressing EGFP-TDP-43 and dnPDI also treated with 1 mM H_2_O_2_ for 1 h, using N2a cells or HEK-293T cells transiently expressing EGFP-TDP-43 and pcDNA3.1 as a control. The above cells were doubly immunostained with the anti-PDI antibody (red) and anti-pTDP-43 antibody (magenta), stained with DAPI (blue), and observed by confocal microscopy (Fig. 5C and fig. S6C). Under pathological stress conditions, EGFP-TDP-43 (green) was partly mislocalized from the nucleus to the cytoplasm of both cell lines and was partly phosphorylated (magenta) (Fig. 5C and fig. S6C). Importantly, we found that under pathological stress conditions, wild-type PDI blocked cytoplasmic mislocalization and phosphorylation of TDP-43 in both cell lines in most fields of view, but dnPDI did not block the cytoplasmic mislocalization and phosphorylation of TDP-43 (Fig. 5C and fig. S6C). We observed that wild-type PDI specifically interacted with TDP-43, including phosphorylated TDP-43 in the cytoplasm of cells under pathological stress conditions (Fig. 5C and fig. S6C). The sarkosyl-insoluble pellets from the same N2a cells as in Fig. 5A were probed with anti-TDP-43 antibody, and the corresponding cell lysates were probed using anti-TDP-43 antibody, anti-HA antibody, and anti-β-actin antibody (Fig. 5D). Treatment of H_2_O_2_ strongly promoted the pathological aggregation of endogenous TDP-43 in N2a cells (Fig. 5D). We then compared the amount of endogenous TDP-43 aggregates (including insoluble full-length TDP-43 and insoluble C-terminal TDP-43 fragment of 35 kDa in Fig. 5D) in N2a cells in the absence and presence of transiently expressed PDI (Fig. 5E). Importantly, we found that under pathological stress conditions, the amount of endogenous TDP-43 aggregates in N2a cells in the presence of wild-type PDI (0.290 ± 0.028, *p* = 0.000091) was significantly lower than that in the absence of PDI (1.45 ± 0.12), but dnPDI only mildly changed the amount of endogenous TDP-43 aggregates in N2a cells (1.04 ± 0.10, *p* = 0.0109) (Fig. 5E). Notably, wild-type PDI blocked the oxidative stress-induced fragmentation and aggregation of endogenous TDP-43 (Fig. 5, D and E).

**Fig. 5.**
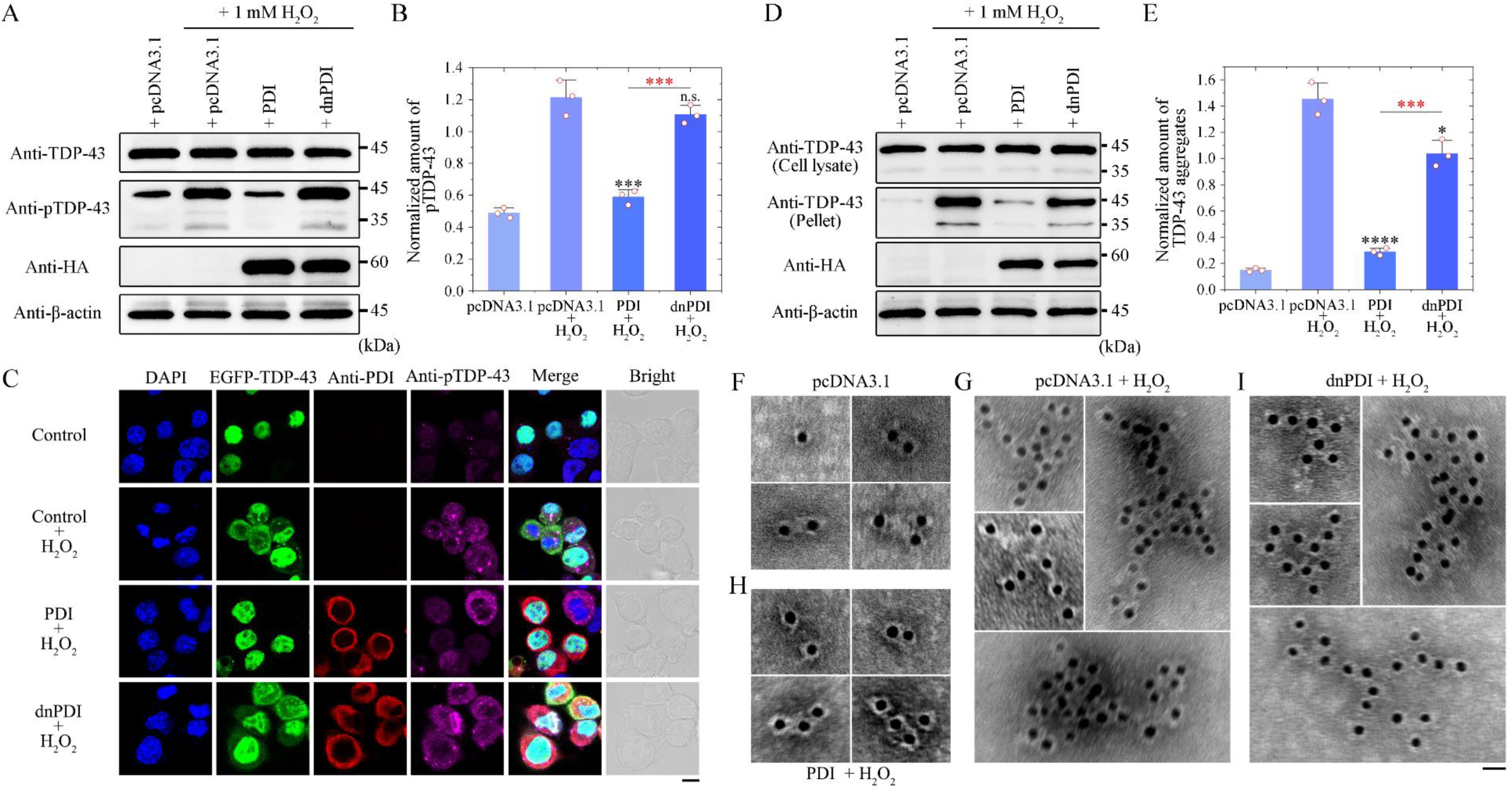
Wild-type PDI blocks pathological phosphorylation and aggregation of TDP-43 induced by H_2_O_2_. (**A**) Western blot for phosphorylated endogenous TDP-43 (pTDP-43) in N2a cells transiently expressing empty vector pcDNA3.1, transiently expressing pcDNA3.1 treated with 1 mM H_2_O_2_ for 1 h, transiently expressing wild-type PDI treated with 1 mM H_2_O_2_ for 1 h, and transiently expressing dnPDI also treated with 1 mM H_2_O_2_ for 1 h. The cell lysates from the aforementioned cells were probed by anti-TDP-43, anti-pTDP-43 (pSer409/410), anti-HA, and anti-β-actin antibodies. (**B**) The normalized amount of pTDP-43 in the aforementioned cells (open red circles shown in scatter plots) was expressed as the mean ± SD (with error bars) of values obtained in three independent experiments. Wild-type PDI + H_2_O_2_, *P* = 0.00083; and dnPDI + H_2_O_2_, *P* = 0.211 or 0.00026. (**C**) Immunofluorescence imaging of N2a cells transiently expressing both EGFP-TDP-43 and pcDNA3.1 (control), transiently expressing both EGFP-TDP-43 and pcDNA3.1 treated with 1 mM H_2_O_2_ for 1 h (control + H_2_O_2_), transiently expressing both EGFP-TDP-43 and wild-type PDI treated with 1 mM H_2_O_2_ for 1 h (wild-type PDI + H_2_O_2_), and transiently expressing both EGFP-TDP-43 and dnPDI also treated with 1 mM H_2_O_2_ for 1 h (dnPDI + H_2_O_2_), using antibodies against PDI (red) and pTDP-43 (magenta) and staining with DAPI (blue). EGFP-TDP-43 (green) was observed in (**C**). Scale bar, 10 μm. (**D**) Western blot for endogenous TDP-43 in the sarkosyl-insoluble pellets (including insoluble full-length TDP-43 and insoluble C-terminal TDP-43 fragment of 35 kDa) and the corresponding cell lysates from the same N2a cells as in (A). β-actin served as the protein loading control. (**E**) The normalized amount of TDP-43 aggregates in the aforementioned cells (open red circles shown in scatter plots) was expressed as the mean ± SD (with error bars) of values obtained in three independent experiments. Wild-type PDI + H_2_O_2_, *P* = 0.000091; and dnPDI + H_2_O_2_, *P* = 0.0109 or 0.00026. Statistical analyses were performed using two-sided Student’s *t*-test. Values of *P* < 0.05 indicate statistically significant differences. The following notation is used throughout: \**P* < 0.05; \*\**P* < 0.01; \*\*\**P* < 0.001; and \*\*\*\**P* < 0.0001 relative to control (pcDNA3.1 + H_2_O_2_ or wild-type PDI + H_2_O_2_). n.s., no significance. (**F** to **I**) Immunogold electron microscopy of TDP-43 aggregates purified from the same N2a cells transiently expressing pcDNA3.1 (F and G), wild-type PDI (H), or dnPDI (I) treated without (F) or with (G−I) 1 mM H_2_O_2_ as in (A), and labeled by gold particles conjugated with anti-TDP-43 antibody. Scale bar, 20 nm.

To ascertain the nature of oxidative stress-induced pathological aggregates of TDP-43 in neuronal cells, we conducted immunogold electron microscopy. The same N2a cells transiently expressing pcDNA3.1 (Fig. 5, F and G), wild-type PDI (Fig. 5H), or dnPDI (Fig. 5I) were treated without (Fig. 5F) or with (Fig. 5, G to I) 1 mM H_2_O_2_ as in Fig. 5A, and labeled by gold particles conjugated with anti-TDP-43 antibody. The amyloid fibrils in the above cell samples were recognized by anti-TDP-43 antibody and decorated with 10-nm gold labels, and the neuronal cells in the presence of wild-type PDI produced much less TDP-43 amyloid fibrils (Fig. 5H) than those in the absence of PDI (Fig. 5G), suggesting that wild-type PDI might function as a disaggregase to reverse ALS’s-linked TDP-43 amyloid fibrils. In sharp contrast, dnPDI did not significantly change the amount of TDP-43 amyloid fibrils (Fig. 5I).

Given that wild-type PDI blocks the pathological phosphorylation and aggregation of TDP-43 in neuronal cells under the oxidative stress condition (Fig. 5), we predicted that wild-type PDI might suppress oxidative stress-induced mitochondrial damage and TDP-43 toxicity. We next used ultrathin section transmission electron microscopy (TEM) (*9*, *52*, *53*) and flow cytometry with annexin V-FITC and propidium iodide (PI) staining (*8*, *52*) to test this hypothesis. To further test the functional relevance of the observations that we made *in vitro* and *in vivo* with TDP-43 LLPS, we used N2a cells stably expressing TDP-43 and transiently expressing empty vector pcDNA3.1, and the stable N2a cells transiently expressing pcDNA3.1 treated with 1 mM H_2_O_2_ for 1 h, transiently expressing wild-type PDI treated with 1 mM H_2_O_2_ for 1 h, and transiently expressing dnPDI also treated with 1 mM H_2_O_2_ for 1 h, using N2a cells transiently expressing pcDNA3.1, transiently expressing pcDNA3.1 treated with 1 mM H_2_O_2_ for 1 h, transiently expressing wild-type PDI treated with 1 mM H_2_O_2_ for 1 h, and transiently expressing dnPDI also treated with 1 mM H_2_O_2_ for 1 h as a control (Fig. 6, A and B). The above cells were detected by flow cytometry with annexin V-FITC and PI staining (Fig. 6, A and B). Treatment of H_2_O_2_ markedly increased toxicity in N2a cells stably expressing TDP-43 compared to wild-type N2a cells (control) (Fig. 6, A and B). Importantly, we found that wild-type PDI remarkably decreased oxidative stress-induced toxicity in N2a cells stably expressing TDP-43 (Fig. 6, A and B). By contrast, wild-type PDI did not remarkably decrease oxidative stress-induced toxicity in wild-type N2a cells (control) and dnPDI did not remarkably decrease oxidative stress-induced toxicity in both wild-type N2a cells and N2a cells stably expressing TDP-43 (Fig. 6, A and B). The morphology of mitochondria in the same N2a cells as in Fig. 6B were then examined by ultrathin section TEM (Fig. 6, C and D). The morphology of normal mitochondria in N2a cells stably expressing TDP-43 and transiently expressing pcDNA3.1, which are highlighted by blue arrows, was tubular or round (Fig. 6, C and D). Treatment of H_2_O_2_ caused severe mitochondrial impairment in N2a cells stably expressing TDP-43 and most of the mitochondria in the cells became swollen and vacuolized, with mitochondria cristae rupturing and disappearance, which are highlighted by red arrows (Fig. 6, C and D). Importantly, we found that wild-type PDI blocked the oxidative stress-induced mitochondrial damage in N2a cells stably expressing TDP-43 and more than half of the mitochondria in the cells were normal in appearance (highlighted by blue arrows) (Fig. 6, C and D). By contrast, dnPDI did not block oxidative stress-induced mitochondrial impairment in N2a cells stably expressing TDP-43 (Fig. 6, C and D). We then compared the number of normal mitochondria in N2a cells stably expressing TDP-43 in the absence and presence of transiently expressed PDI (Fig. 6E). Importantly, a significantly higher number of normal mitochondria was observed in N2a cells stably expressing TDP-43 in the presence of wild-type PDI ((58.0 ± 5.5)%, *p* = 8.5 × 10^-44^) than that in the cell line in the absence of PDI ((13.4 ± 2.7)%). A similar conclusion was reached in in HEK-293T cells stably expressing TDP-43 (fig. S7, A to C). Therefore, wild-type PDI significantly decreases mitochondrial damage caused by TDP-43 aggregation and induced by oxidative stress, possibly via specific interactions with TDP-43.

**Fig. 6.**
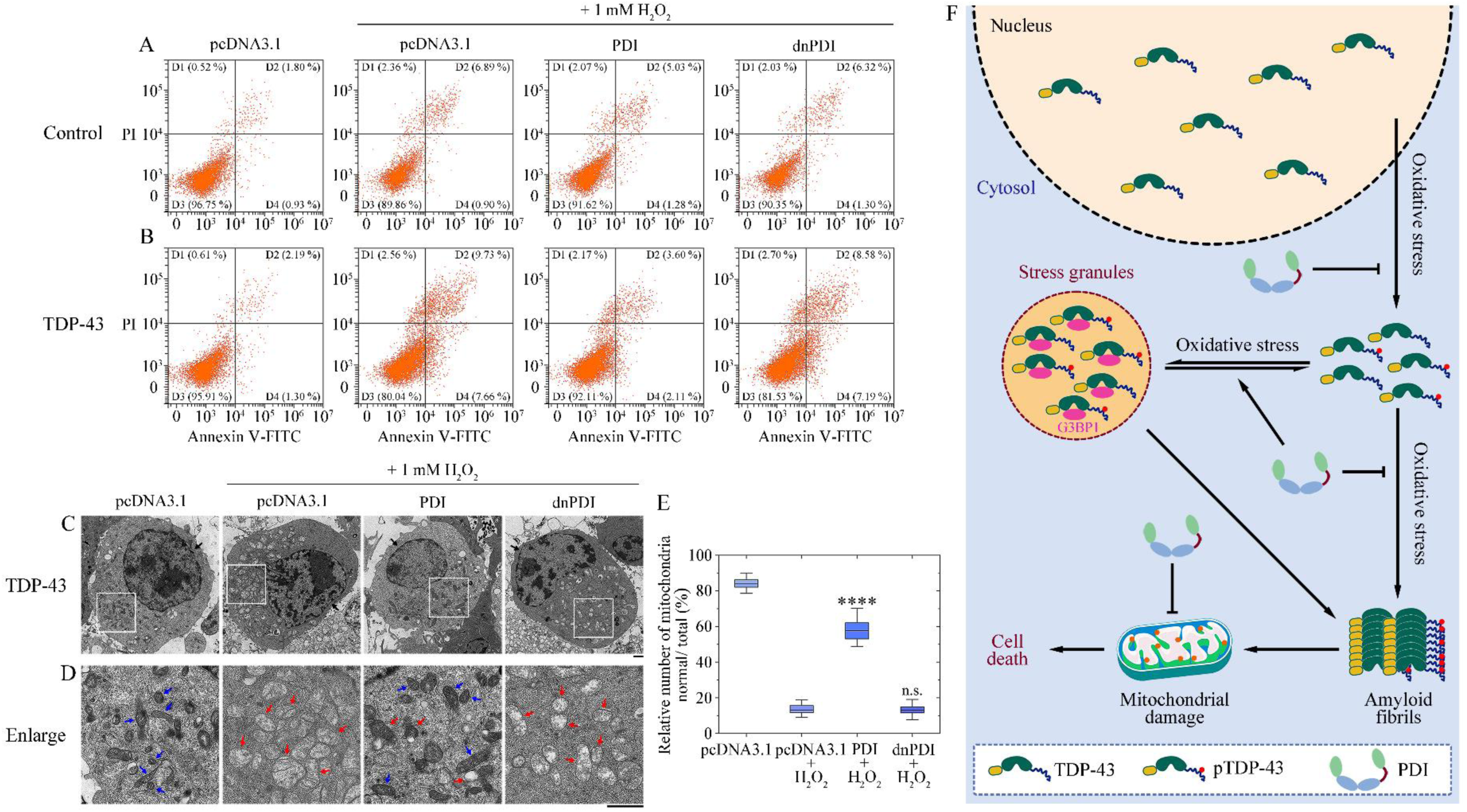
Wild-type PDI disassembles stress granules, blocks cytoplasmic mislocalization and aggregation of TDP-43, and suppress mitochondrial damage and TDP-43 toxicity. (**A**) Wild-type PDI did not remarkably decrease H_2_O_2_-induced toxicity in wild-type N2a cells (control). Cell viability: 96.8% for N2a cells transiently expressing empty vector pcDNA3.1; 89.9% for N2a cells transiently expressing pcDNA3.1 treated with 1 mM H_2_O_2_ for 1 h; 91.6% for N2a cells transiently expressing wild-type PDI treated with 1 mM H_2_O_2_ for 1 h; and 90.4% for N2a cells transiently expressing dnPDI also treated with 1 mM H_2_O_2_ for 1 h. (**B**) Wild-type PDI remarkably decreased H_2_O_2_-induced toxicity in N2a cells stably expressing TDP-43. Cell viability: 95.9% for stable N2a cells transiently expressing pcDNA3.1; 80.0% for stable N2a cells transiently expressing pcDNA3.1 treated with 1 mM H_2_O_2_ for 1 h; 92.1% for stable N2a cells transiently expressing wild-type PDI treated with 1 mM H_2_O_2_ for 1 h; and 81.5% for stable N2a cells transiently expressing dnPDI also treated with 1 mM H_2_O_2_ for 1 h. (**C** and **D**) Wild-type PDI blocked H_2_O_2_-induced mitochondrial damage in N2a cells stably expressing TDP-43. TEM imaging of stable N2a cells transiently expressing pcDNA3.1, transiently expressing pcDNA3.1 treated with 1 mM H_2_O_2_ for 1 h, transiently expressing wild-type PDI treated with 1 mM H_2_O_2_ for 1 h, and transiently expressing dnPDI also treated with 1 mM H_2_O_2_ for 1 h (C). The enlarged regions (D) show 14-fold enlarged images from (C) and display the detailed structures of mitochondria in cells. Nuclei and normal mitochondria in N2a cells are highlighted using black arrows and blue arrows, respectively. Abnormal mitochondria in cells with mitochondrial vacuolization are highlighted by red arrows. Samples were negatively stained using 2% uranyl acetate and lead citrate. Scale bars, 1 μm. (**E**) Wild-type PDI significantly decreased H_2_O_2_-induced mitochondrial damage in N2a cells stably expressing TDP-43. Box plots analyzing the relative number of mitochondria (normal/total) in the aforementioned cells and showing the quantification of TEM images in *n* = 30 cells examined over 3 independent experiments. The experiments were done blind and average 42 to 57 mitochondria were present in each cell. The boxes (blue) extend from the 25th to 75th percentile (quantiles 1 and 3). Minima, maxima, centre, and bounds of box represent quantile 1 minus 1.5 × interquartile range, quantile 3 plus 1.5 × interquartile range, median (black line), and quantiles 1 and 3, respectively. Wild-type PDI + H_2_O_2_, *P* = 8.5 × 10^-44^; and dnPDI + H_2_O_2_, *P* = 0.973. Statistical analyses were performed using two-sided Student’s *t*-test. Values of *P* < 0.05 indicate statistically significant differences. The following notation is used throughout: \**P* < 0.05; \*\**P* < 0.01; \*\*\**P* < 0.001; and \*\*\*\**P* < 0.0001 relative to control (pcDNA3.1 + H_2_O_2_). n.s., no significance. (**F**) A hypothetical model shows how wild-type PDI disassembles stress granules and blocks cytoplasmic mislocalization and aggregation of TDP-43. Under oxidative stress conditions, TDP-43 (N-terminal domain, yellow; C-terminal domain, ultramarine; and two RNA recognition motifs, dark green) is partly mislocalized from the nucleus (light yellow) to the cytosol (pale blue) and is hyperphosphorylated (red), but wild-type PDI (domains a and a’, green; domains b and b’, blue; and the x linker, dark red) blocks cytoplasmic mislocalization and phosphorylation of TDP-43 (pTDP-43). Then, TDP-43 selectively interacts with G3BP1 (magenta ellipsoid) to assemble stress granules (orange balls), but wild-type PDI disassembles stress granules via specific interactions with TDP-43. Finally, TDP-43 and stress granules are assembled into amyloid fibrils, resulting in mitochondrial damage and neuronal cell death in patients with ALS and AD, but wild-type PDI blocks pathological aggregation of TDP-43 and suppress mitochondrial damage and TDP-43 toxicity.

Altogether, these data demonstrate that TDP-43 recruits wild-type PDI into its phase-separated condensate in neuronal cells under pathological stress condition, which in turn blocks the oxidative stress-induced pathological phosphorylation and aggregation of TDP-43 to form much less aggregates and to suppress mitochondrial damage and TDP-43 toxicity in ALS.

## DISCUSSION

TDP-43, a ubiquitous protein encoded by the *TARDBP* gene, has important functions in normal cellular physiology, such as precursor mRNA (pre-mRNA) maturation (*1*, *61*). By contrast, TDP-43 dysfunction, characterized by cytoplasmic mislocalization and aggregation of the protein, is not only responsible for neuronal cell death in ALS, FTLD-TDP, and other TDP-43 proteinopathies but also observed in the brain of some AD patients (*1*−*6*, *10*−*18*, *62*). In this study, we detected a much higher level of cytoplasmic TDP-43 in the brain and spinal cord from two ALS patients and in the brain from six AD patients than in the brain from three healthy controls, one control with Castleman disease, and one control with PSP. Accumulating pieces of evidence point to a crucial role of oxidative stress in ALS pathogenesis (*46*, *47*, *55*, *56*). We showed that oxidative stress caused TDP-43 to mislocalize and accumulate in the cytoplasm of neuronal cells, promoted subsequent pathological phosphorylation, fragmentation and aggregation of TDP-43, and increased TDP-43 toxicity in ALS.

Several in vivo co-IP studies have described physical interactions between TDP-43 and different classes of proteins in neurons and neuronal cells, among which amyloid-β, Rho guanine nucleotide exchange factor, Hsp90 and its co-chaperone Sti1, histone deacetylase 1 and Hsp70, and Grp78, are highlighted as potential TDP-43 partners (*14*, *63*−*66*). Given that PDI is partly mislocalized from the ER to the cytoplasm under ER stress condition (this work, *22*, *24*−*29*, *67*) and wild-type PDI plays important roles in preventing protein aggregation in ALS pathogenesis (this work, *22*−*24*), we predicted that under pathological stress conditions, wild-type PDI might physically interact with TDP-43 in the cytoplasm to modulate phosphorylation and aggregation of TDP-43. In this study, we found wild-type PDI associated with endogenous or overexpressed TDP-43 in the cytoplasm. We showed that cytoplasmic TDP-43 accumulated and colocalized with PDI in not only ALS brain and spinal cord samples but also AD brain samples, and that overexpressed TDP-43 colocalized with wild-type PDI in the cytoplasm of neuronal cells upon stress. Wild-type PDI physically interacts with TDP-43 in the cytoplasm to block pathological phosphorylation, fragmentation, and aggregation of TDP-43 and to suppress TDP-43 toxicity in ALS. Therefore, PDI regulates pathological functions of cytoplasmic TDP-43 in neurons via multiple mechanisms, the first being physical interaction with TDP-43.

In this work, we report that TDP-43, a multi-facet protein existing in cytoplasmic form under pathological stress conditions, exhibits disparate propensities to phase separate with PDI. We show that TDP-43 undergoes LLPS in vitro and within stressed cells. TDP-43 condensates selectively recruit wild-type PDI, which in turn strongly slows down the LLPS of TDP-43 in vitro and within stressed cells. TDP-43 also recruits and concentrates an abnormal form of PDI into phase-separated condensates (puncta) in the cytosol of stressed cells, which in turn mediates TDP-43 condensation, rendering the resulting liquid droplets fibril-like. Overall, our results show that TDP-43 undergoes PDI-mediated LLPS in vitro and within stressed cells. The recruitment of wild-type PDI into TDP-43 condensates weakens the interaction between TDP-43 and G3BP1 under pathological stress conditions, results in the disassembly of stress granules in neuronal cells, and blocks pathological aggregation of TDP-43. It has been reported that *in vivo* LLPS of TDP-43 is modulated by two types of molecular chaperones (Hsp70 and HSPB1) (*37*, *39*). Intriguingly, TDP-43 condensates selectively recruit HSPB1 in cells upon stress, which in turn inhibits TDP-43 assembly into fibrils, and the combined activities of HSPB1, BAG2, and HSPA1A facilitate the disassembly of TDP-43 condensates within stressed cells (*39*). It should be mentioned that experimental disruption of G3BP1 dimerization, ubiquitination of G3BP1, and two G3BP inhibitors all result in the disassembly of stress granules in cells (*57*, *58*, *60*). Thus, the specific properties of proteins included in TDP-43 condensates influence the liquidness and organization of TDP-43 condensates (this work, *37*, *39*), and targeting of the core protein G3BP1 weakens the stress granule–specific interactions, resulting in the disassembly of stress granules in cells (this work, *58*, *60*).

In summary, our results describe a model to underpin molecular hypotheses of how wild-type PDI disassembles stress granules and blocks cytoplasmic mislocalization and aggregation of TDP-43 in ALS (Fig. 6F). We report interesting findings on the roles of cytoplasmic TDP-43, LLPS composed of cytoplasmic TDP-43 and PDIs, and certain PDIs on ALS and other neurodegenerative diseases associated with accumulation or overexpression of cytoplasmic TDP-43 (TDP-43 proteinopathies). Importantly, we show that under pathological stress condition, TDP-43, the major component of pathological inclusions in most patients with ALS and FTLD-TDP and in up to 57% of patients with AD (*1*−*6*, *10*−*18*), is partly mislocalized from the nucleus to the cytoplasm of neuronal cells, is hyperphosphorylated, and selectively interacts with wild-type PDI; this interaction in turn blocks cytoplasmic mislocalization and phosphorylation of TDP-43 (Fig. 6F). Then, TDP-43 selectively interacts with G3BP1 to assemble stress granules, but wild-type PDI disassembles stress granules via specific interactions with TDP-43 (Fig. 6F). In response to oxidative stress, TDP-43 recruits and interacts with proteins of the ER and endosomal-extracellular transport pathways (*66*), such as wild-type PDI (this work). Finally, in the presence of abnormal forms of PDI, PDI loses its activity, TDP-43 and stress granules are assembled into amyloid fibrils, resulting in mitochondrial damage and neuronal cell death in ALS patients and some Alzheimer’s patients, but wild-type PDI blocks pathological aggregation of TDP-43 and suppresses mitochondrial impairment and TDP-43 toxicity (Fig. 6F). Therefore, accumulation or overexpression of cytoplasmic TDP-43 is toxic to neurons via enhancing pathological phosphorylation, fragmentation, and aggregation of TDP-43 under oxidative stress conditions, and that wild-type PDI suppresses the TDP-43 toxicity. These results provide insights on the regulation of PDI on pathological functions of cytoplasmic TDP-43 via LLPS of the protein inhibited by wild-type PDI, but not abnormal forms of PDI, which has important implications in ALS etiology. The observation of many colocalized dots of TDP-43 and PDI in the cytoplasm of a group of ALS patients and AD patients provides clinical relevance. The selective interaction of wild-type PDI with TDP-43 upon stress will be valuable to understanding the functional basis underlying LLPS of proteins and inspiring future research on protein condensation diseases caused by abnormal liquid-like or solid-like states of proteins (*68*) and regulated by protein interaction partners (*37*, *39*, *69*).

## MATERIALS AND METHODS

### Ethics statement

The study complies with all relevant ethical regulations. The study is based on analyses of brain and spinal cord samples from two patients with amyotrophic lateral sclerosis, and brain samples from six patients with Alzheimer’s disease and from three healthy individuals, one patient with Castleman disease, and one patient with progressive supranuclear palsy (controls). The patient characteristics are described in Table 1. Post-mortem human brain tissue samples were obtained from the National Human Brain Bank for Development and Function, Institute of Basic Medical Sciences Chinese Academy of Medical Sciences, School of Basic Medicine Peking Union Medical College (Case 1 and Control 4) and the Department of Neurology, Xiangya Hospital, Central South University (Cases 3 to 8, Controls 1 to 3, and Control 5), and the human post-mortem spinal cord samples were obtained from the Department of Neurology, Peking University Third Hospital (Case 2). Their use in this study was approved by the ethical review processes at each institution, including ethical approval from relevant authorities at Wuhan University. Informed consent was obtained from the patients or their next of kin. We have obtained consent to publish information that identifies individuals (including three or more indirect identifiers such as exact age, sex, and medical centre the study participants attended or rare diagnosis). The biochemical work at Wuhan University was conducted based on a permission from the Wuhan University Ethics Committee (WAEF-2022-0073).

### Double immunofluorescence staining of pathological samples

Sections of the precentral gyrus (Case 1 and Control 4), spinal cord (Case 2), and hippocampus and temporal lobe (Cases 3 to 8, Controls 1 to 3, and Control 5) were formalin-fixed and paraffin-embedded for *in situ* staining. The paraffin-embedded sections of 5 μm were cut, deparaffinized in xylene, and rehydrated in a descending series of ethanol. Antigen retrieval was carried out using an Improved Citrate Antigen Retrieval Solution (Beyotime, P0083) at 95-100 °C for 20 min, and cooled to room temperature. After being washed three times with PBS for 3 min, sections were marked with hydrophobic circles using an immunohistochemical pen. After being permeabilized with cold 0.3% Triton X-100 in PBS at room temperature for 15 min, sections were blocked using 5% BSA at room temperature for 1 h, followed by overnight incubation with the primary antibodies, diluted in PBS containing 5% BSA in a humidified chamber at 4 °C. After three times washes with PBS for 5 min, fluorescence-conjugated secondary antibodies were applied to the cover slip and incubated in a dark room for 1 h. DAPI was then applied at the proper dilution; after being washed three times with PBS for 5 min and mounted with antifade mounting medium (Beyotime, P0126), cells were subjected to a Leica TCS SP8 laser scanning confocal microscope (Wetzlar, Germany). The following primary antibodies were used: rabbit anti-TDP-43 polyclonal antibody (Proteintech, 10782-2-AP, 1:200), mouse anti-P4HB monoclonal antibody (anti-PDI antibody, Abcam, ab2792, 1:200), Alexa Fluor 488-labeled goat anti-rabbit IgG (H+L) (Beyotime, A0423, 1:500), and Alexa Fluor 555-labeled donkey anti-mouse IgG (H+L) (Beyotime, A0460, 1:500).

### H&E staining of pathological samples

H&E staining of the paraffin brain and spinal cord sections was conducted by following the manufacturer’s instructions (Beyotime, C0105S). In brief, slices were deparaffinized in xylene, and rehydrated in a descending series of ethanol. The slices were stained with hematoxylin for 15 min, and then washed with running tap water for 10 min, followed by incubation with hydrochloric acid alcohol solution (Beyotime, C0163L) for 10 s. After being washed with running tap water for 10 min, the slices were stained with eosin for 1 min and dehydrated in an incremental series of ethanol. After being treated twice with xylene for 5 min, the slices were mounted with neutral balsam (Biosharp, BL704A) and imaged using a Leica Aperio Versa slide scanner (Wetzlar, Germany).

### Protein expression and purification

A plasmid-encoding, human full-length TDP-43 was a kind gift from Dr. H.-N. Du (College of Life Sciences, Wuhan University). Recombinant, full-length wild-type human TDP-43 (residues 1−414) containing a tag of six histidine residues (polyhistidine tag) at its C-terminal domain was constructed into a prokaryotic expression vector pET22b and expressed in *E. coli* BL21 (DE3) cells (Novagen, Merck, Darmstadt, Germany). TDP-43 protein was purified to homogeneity by nickel affinity chromatography as described by Vega et al. (*70*). After purification, the refolded TDP-43 protein was dialyzed against 50 mM Tris-HCl buffer (pH 8.0) containing 150 mM NaCl four times to remove the detergents and salts, concentrated, and then centrifuged at 17,000 *g* for 30 min at 4 °C to remove large aggregates. The supernatant of the refolded TDP-43 protein was freshly used or stored at −80 °C. SDS-PAGE was used to confirm that the purified human TDP-43 protein was single species. We used a BCA protein assay kit (Beyotime, P0012) to determine the concentration of human TDP-43 protein.

A plasmid-encoding, human full-length PDI was a kind gift from Dr. L. W. Ruddock (Faculty of Biochemistry and Molecular Medicine, University of Oulu). The gene for PDI 1-491 was constructed in a prokaryotic expression vector pET23, and a PDI mutant dnPDI was constructed by site-directed mutagenesis using a wild-type PDI template; the primers are shown in table S1. All PDI plasmids were transformed into *Escherichia coli*. Recombinant full-length wild-type human PDI (residues 1−491) and its variant dnPDI were expressed from the vector pET23 in *E. coli* BL21 (DE3) Codon plus-RIL cells (Novagen, Merck, Darmstadt, Germany). PDI proteins were purified to homogeneity by nickel affinity chromatography as described by Wang et al. (*71*). The eluted fractions were dialyzed against 50 mM Tris-HCl buffer (pH 8.0) containing 150 mM NaCl twice to remove EDTA. The PDI proteins were freshly used or stored at - 80 °C. SDS-PAGE was used to confirm that the purified human PDI proteins were single species. We used a UV-2550 Probe spectrophotometer (Shimadzu, Kyoto, Japan) to determine the concentrations of wild-type human PDI and dnPDI, using their absorbances at 214 nm with a standard calibration curve drawn by BSA.

### Characterization of SNO-PDI by fluorescence spectroscopy

S-nitrosoglutathione (GSNO) was synthesized as described in detail by Jones et al. (*72*). To produce SNO-PDI, wild-type PDI and GSNO were cultured at a molar ratio of 1:100 and incubated for 30 min at room temperature, then concentrated the protein, and washed with 50 mM Tris-HCl buffer (pH 8.0) containing 150 mM NaCl to remove residual GSNO, using centrifugal filters. SNO-PDI was then assessed by the release of NO, causing the conversion of 2,3-diaminonaphthalene (DAN) into the fluorescent compound 2,3-naphthyltriazole (NAT) (*21*, *73*, *74*). In brief, Tris-HCl buffer (pH 8.0) (control) and Tris-HCl buffer (pH 8.0) containing wild-type PDI or the purified SNO-PDI were incubated with a solution containing 100 μM DAN and 100 μM CuSO_4_ in the dark at room temperature for 30 min. 1 M NaOH was then added into each tube to stop the reaction. The NO-dependent S-nitrosylation of DAN to yield its highly fluorescent S-nitrosylated derivative, NAT, was quantified by measuring the fluorescence of each sample using an excitation wavelength at 375 nm and an emission wavelength at 450 nm. In other words, the fluorescence intensity of NAT in the above samples was measured using excitation at 375 nm and emission at 450 nm.

### Liquid-droplet formation

The freshly bacterial-purified wild-type human TDP-43 was incubated with TAMRA (red fluorescence, excitation at 561 nm) at a TDP-43: TAMRA molar ratio of 1:3 at room temperature for 1 h; the freshly bacterial-purified wild-type human PDI and two abnormal forms of PDI (dnPDI and SNO-PDI), along with BSA, a negative control, were incubated with FITC (green fluorescence, excitation at 488 nm) at a PDI/BSA: FITC mass ratio of 1:50-80 in the dark at 4 °C for at least 8 h. These labeled proteins were filtered, concentrated in a centrifugal filter (Millipore) and diluted in 50 mM Tris-HCl buffer (pH 8.0) containing 150 mM NaCl. 10 μM TDP-43 labeled by TAMRA was incubated with Tris-HCl buffer (pH 8.0) containing 150 mM NaCl and 10% (w/v) PEG 3350 or incubated with the same buffer further containing 10 μM wild-type PDI, dnPDI, or SNO-PDI labeled by FITC on ice to induce LLPS, and FITC-labeled BSA as a control. TDP-43 de-mixed droplets (red) coacervated with PDI labeled by FITC (green) were observed by a Leica TCS SP8 laser scanning confocal microscope (Wetzlar, Germany), with excitation at 561 nm and 488 nm, respectively. In total, 2.5, 5, 7.5, or 10 μM TDP-43 labeled by TAMRA was incubated with Tris-HCl buffer (pH 8.0) containing 150 mM NaCl and 10% PEG 3350 or incubated with the same buffer further containing 10 μM wild-type PDI, dnPDI, or SNO-PDI on ice to induce LLPS. Liquid droplets of TDP-43 were observed by a Leica TCS SP8 laser scanning confocal microscope (Wetzlar, Germany) with excitation at 561 nm. All phase separation experiments were performed at least three times and were pretty reproducible.

### FRAP

In total, 10 μM wild-type human TDP-43 labeled by TAMRA was incubated with Tris-HCl buffer (pH 8.0) containing 150 mM NaCl and 10% (w/v) PEG 3350 or incubated with the same buffer further containing 10 μM wild-type human PDI, dnPDI, or SNO-PDI on ice to induce LLPS. Liquid droplets of TDP-43 were observed by a Zeiss LSM 880 laser scanning confocal microscope equipped with an Airyscan detector (Carl Zeiss, Germany), with excitation at 561 nm. We chose TDP-43 droplets with sizes of ∼ 5 μm for FRAP experiments. For each droplet, a square was bleached at 80% transmission for one time, and postbleaching time-lapse images were collected (250 frames, 1000 ms per frame). Images were analyzed using Zen (LSM 880 confocal microscope manufacturer’s software). All FRAP experiments were repeated three times and the results were reproducible.

### Turbidity assays

In total, 0.50, 1.0, 2.0, 3.0, 4.0, 5.0, 6.0, 7.0, 8.0, 9.0, and 10 μM wild-type human TDP-43 was incubated with Tris-HCl buffer (pH 8.0) containing 150 mM NaCl and 10% PEG 3350 or incubated with the same buffer further containing 10 μM wild-type human PDI, dnPDI, or SNO-PDI on ice to induce LLPS. The turbidity of TDP-43 condensates was measured at 600 nm and 25 °C using a NanoDrop OneC Microvolume UV-Vis Spectrophotometer (Thermo Fisher Scientific). All turbidity assays were repeated at least three times and were pretty reproducible.

### Cell culture and transfection

N2a neuroblastoma cells (catalog number GDC0162) and HEK-293T cells (catalog number GDC0187) were obtained from China Center for Type Culture Collection (CCTCC, Wuhan, China) and cultured in minimum essential media and in Dulbecco’s modified Eagle’s medium (Gibco, Invitrogen), respectively, supplemented with 10% (v/v) fetal bovine serum (Gibco), 100 U/ml streptomycin, and 100 U/ml penicillin in 5% CO_2_ at 37 °C. Transient transfections were performed using Lipofectamine 2000 (Invitrogen) according to the manufacturer’s protocol. N2a and HEK-293T cells were transiently transfected with phage-TDP-43, phage-EGFP-TDP-43, pEGFP-TDP-43-EGFP_1-172_, pcDNA3.1-EGFP_155-238_-PDI, pcDNA3.1-EGFP_155-238_-dnPDI, pEGFP-EGFP_1-172_, pcDNA3.1-EGFP_155-238_, empty vector pcDNA3.1, pcDNA3.1-HA-PDI, and pcDNA3.1-HA-dnPDI as indicated. In brief, after one-day culture, 2 μg plasmid and 5 μl Lipofectamine 2000 (Invitrogen) were diluted into 200 μl Opti-MEM (Gibco) in each well of a 6-well plate cells, the transfection solution was incubated in 5% CO_2_ at 37 ℃ for 4–6 h, and then cultured in proper cell culture medium for 36-48 h. N2a and HEK-293T cell lines stably expressing wild-type human TDP-43 were constructed with a lentiviral vector construction system (phage-puro). The target DNA fragments were inserted into the lentiviral vector, and the plasmids containing target DNA, pVSVG, and p976 were packaged in HEK-293T cells at a ratio of 2:1:1 by Lipofectamine 2000 (Invitrogen). The ratio of liposome to DNA was 2:1. After 48 h of transfection, the viruses were harvested and filtered, and then N2a cells and HEK-293T cells were infected with the packaged lentivirus twice for 12 h each with a 12-h interval. In order to establish the stable cell lines, puromycin was used to screen overexpressed cells. The expression of each protein was detected by Western blot.

### Western blotting

For analysis by western blotting, N2a and HEK-293T cells grown in a 6-well plate for the day were transfected as indicated. Cells were washed twice with ice-cold PBS and lysed in 300 μl (per well) cell lysis buffer containing 1 × protease inhibitor cocktail (Target Mol). The amount of loaded protein was normalized using a BCA Protein Quantification kit (Beyotime, P0012). The cell lysates were boiled in SDS-PAGE loading buffer for 10 min and then subjected to SDS-PAGE and probed with the following specific antibodies: mouse anti-TDP-43 monoclonal antibody (Abcam, ab57105,1:1000), rabbit anti-TDP-43 polyclonal antibody (Proteintech, 10782-2-AP, 1:2000), rabbit anti-HA tag polyclonal antibody-ChIP Grade (Abcam, ab9110, 1:1000), mouse anti-HA tag monoclonal antibody (Proteintech, 66006-2-Ig, 1:5000), rabbit anti-Histone-H3 polyclonal antibody (Proteintech, 17168-1-AP, 1:1000), mouse anti-G3BP1 monoclonal antibody (Proteintech, 66486-1-Ig, 1:5000), rabbit anti-phospho-TDP-43 (pS409/410) polyclonal antibody (Proteintech, 22309-1-AP, 1:1000), mouse anti-β-actin monoclonal antibody (Proteintech, 66009-1-Ig, 1:5000), HRP-labeled goat anti-rabbit IgG (H+L) (Beyotime, A0208, 1:5000), and HRP-labeled goat anti-mouse IgG (H+L) (Beyotime, A0216, 1:5000).

### Coimmunoprecipitation

To confirm the interaction between TDP-43 and PDI, HEK-293T cells were transiently co-transfected with phage-TDP-43 and pcDNA3.1-HA-PDI or pcDNA3.1-HA-dnPDI according to the manufacturer’s instructions. After one-day culture and 1-h treatment with 1 mM H_2_O_2_ at 37 ℃, cells were harvested by centrifugation at 100 *g* at 4 °C for 5 min and ruptured on ice for 20 min with cell lysis buffer containing 1 × protease inhibitor cocktail (Target Mol). The cell lysates were then centrifuged at 17,000 *g* at 4 °C for 20 min to remove cell debris. An input of collected supernatant was set aside for subsequent analysis. Each sample was divided and immunoprecipitated with 1 μl of 1.0 mg/ml mouse anti-TDP-43 antibody or mouse IgG on a rotator overnight at 4 ℃. Complexes were pulled down by incubation with Protein G Agarose beads (20 μl, Fast Flow for IP) (Beyotime) on a rotator at 4 ℃ for 4 h. Beads were washed five times with PBS, were boiled in SDS-PAGE loading buffer for 10 min and then subjected to SDS-PAGE, and then probed with Western blot using rabbit anti-HA antibody. Nonspecific IgG was served as a negative control in immunoprecipitation. Another reserved cell lysates were boiled in SDS-PAGE loading buffer, probed with Western blot using anti-HA antibody, anti-TDP-43 antibody, and anti-β-actin antibody, respectively, and served as the input controls, which represented the total PDI or dnPDI content, the total TDP-43 content, and the total protein content in cell lysates, respectively. The following antibodies were used: mouse anti-TDP-43 monoclonal antibody (Abcam, ab57105,1:1000), rabbit anti-HA tag polyclonal antibody-ChIP Grade (Abcam, ab9110, 1:1000), mouse anti-β-actin monoclonal antibody (Proteintech, 66009-1-Ig, 1:5000), HRP-labeled goat anti-rabbit IgG (H+L) (Beyotime, A0208, 1:5000), and HRP-labeled goat anti-mouse IgG (H+L) (Beyotime, A0216, 1:5000).

To confirm the disassembly of stress granules, N2a cells were transiently transfected with empty vector pcDNA3.1, pcDNA3.1-HA-PDI, or pcDNA3.1-HA-dnPDI according to the manufacturer’s instructions. After one-day culture and 1-h treatment with 1 mM H_2_O_2_ or PBS at 37 ℃, cells were harvested by centrifugation at 100 *g* at 4 °C for 5 min and ruptured on ice for 20 min with cell lysis buffer containing 1 × protease inhibitor cocktail (Target Mol). The cell lysates were then centrifuged at 17,000 *g* at 4 °C for 20 min to remove cell debris. An input of collected supernatant was set aside for subsequent analysis. Each sample was divided and immunoprecipitated with 1 μl of 1.0 mg/ml rabbit anti-TDP-43 antibody or rabbit IgG on a rotator overnight at 4 ℃. Complexes were pulled down by incubation with Protein G Agarose beads (20 μl, Fast Flow for IP) (Beyotime) on a rotator at 4 ℃ for 4 h. Beads were washed five times with PBS, were boiled in SDS-PAGE loading buffer for 10 min and then subjected to SDS-PAGE, and then probed with Western blot using mouse anti-G3BP1 antibody. Nonspecific IgG was served as a negative control in immunoprecipitation. Another reserved cell lysates were boiled in SDS-PAGE loading buffer, probed with Western blot using anti-G3BP1 antibody, anti-HA antibody, anti-TDP-43 antibody, and anti-β-actin antibody, respectively, and served as the input controls, which represented the total G3BP1 content, the total PDI or dnPDI content, the total TDP-43 content, and the total protein content in cell lysates, respectively. The following antibodies were used: rabbit anti-TDP-43 polyclonal antibody (Proteintech, 10782-2-AP, 1:2000), mouse anti-HA tag monoclonal antibody (Proteintech, 66006-2-Ig, 1:5000), mouse anti-G3BP1 monoclonal antibody (Proteintech, 66486-1-Ig, 1:5000), mouse anti-β-actin monoclonal antibody (Proteintech, 66009-1-Ig, 1:5000), HRP-labeled goat anti-rabbit IgG (H+L) (Beyotime, A0208, 1:5000), and HRP-labeled goat anti-mouse IgG (H+L) (Beyotime, A0216, 1:5000).

### Bimolecular fluorescence complementation assay

The human *TARDBP* gene was cloned into the pEGFP-N1 vector, the fusion gene was truncated at amino acid 172 of EGFP using a stop codon, and the pEGFP-TDP-43-EGFP_1-172_ vector was then obtained. Besides, the human *P4HB* gene was cloned into the pcDNA3.1 vector, the fused HA-tag and EGFP_155-238_ genes were inserted into the *P4HB* genes between the signal peptide and the amino acid coding sequence, and HA-tagged pcDNA3.1-EGFP_155-238_-PDI vector was then obtained. pcDNA3.1-EGFP_155-238_-dnPDI vector was obtained by mutated the four cysteines (Cys53, Cys56, Cys397, and Cys400) of pcDNA3.1-EGFP_155-238_-PDI to Ala. The fusion expression vectors were confirmed by DNA sequencing, and the expression of fusion proteins were detected by Western blot. One day before transfection, HEK-293T cells were plated on glass-bottom culture dishes. Cells were then transiently co-transfected with pEGFP-TDP-43-EGFP_1-172_ and pcDNA3.1-EGFP_155-238_-PDI or pcDNA3.1-EGFP_155-238_-dnPDI according to the manufacturer’s instructions. After one-day culture, living cells were stained with Hoechst 33342 (blue), and EGFP (green) and nuclei (blue) were observed by a Leica TCS SP8 laser scanning confocal microscope (Wetzlar, Germany), with excitation at 405 nm and 488 nm, respectively. Confocal microscopy was also used to visualize the fluorescence of the following control cells: living HEK-293T cells transiently expressing both EGFP-TDP-43-EGFP_1-172_ and pcDNA3.1-EGFP_155-238_ constructs, and transiently expressing both EGFP-EGFP_1-172_ and pcDNA3.1-EGFP_155-238_-PDI, pcDNA3.1-EGFP_155-238_-dnPDI, or pcDNA3.1-EGFP_155-238_ constructs.

### Nuclear and cytoplasmic protein fractionation

N2a and HEK-293T cells grown in a 6-well plate for the day were transfected as indicated. Cells were washed three times with ice-cold PBS and lysed in 300 μl (per well) cell lysis buffer containing 1 × protease inhibitor cocktail (Target Mol). The nuclear and cytoplasmic proteins were isolated from cells using Nuclear and Cytoplasmic Protein Extraction Kit (Beyotime, P0027) according to the manufacturer’s instructions. The amount of loaded protein was normalized using a BCA Protein Quantification kit (Beyotime, P0012) and probed with Western blot. The following antibodies were used: rabbit anti-TDP-43 polyclonal antibody (Proteintech, 10782-2-AP, 1:2000), mouse anti-HA tag monoclonal antibody (Proteintech, 66006-2-Ig, 1:5000), rabbit anti-Histone-H3 polyclonal antibody (Proteintech, 17168-1-AP, 1:1000), mouse anti-β-actin monoclonal antibody (Proteintech, 66009-1-Ig, 1:5000), HRP-labeled goat anti-rabbit IgG (H+L) (Beyotime, A0208, 1:5000), and HRP-labeled goat anti-mouse IgG (H+L) (Beyotime, A0216, 1:5000).

### Immunofluorescence staining

N2a and HEK-293T cells plated on coverslips in a 12-well plate for the day were transfected as indicated. Cells were washed three times with PBS, fixed with 4% (w/v) paraformaldehyde in PBS for 30 min, and then permeabilized with 0.25% Triton X-100 in PBS at room temperature for 5 min. Cells were then blocked with 5% BSA in PBS at 37 ℃ for 30 min, and incubated with appropriate dilution ratio of primary antibodies at 37 ℃ for 2 h. After three times washes with PBS for 5 min, fluorescence-conjugated secondary antibodies at appropriate dilution ratios were applied to the samples and incubated in a dark room for 45 min. DAPI was then applied at the proper dilution; after being washed three times with PBS for 5 min and mounted with antifade mounting medium (Beyotime, P0126), cells were subjected to a Leica TCS SP8 laser scanning confocal microscope (Wetzlar, Germany). The following antibodies were used: mouse anti-P4HB monoclonal antibody (anti-PDI antibody, Abcam, ab2792, 1:500), rabbit anti-G3BP1 polyclonal antibody (Proteintech, 13057-2-AP, 1:500), rabbit anti-pTDP-43 (pSer409/410) polyclonal antibody (Cosmo, CAC-TIP-PTD-P07, 1:1000), Alexa Fluor 555-labeled donkey anti-mouse IgG (H + L) (Beyotime, A0460, 1:1000), and Alexa Fluor 647-labeled goat anti-rabbit IgG (H+L) (Beyotime, A0468, 1:1000).

### Sarkosyl-insoluble western blotting

Sarkosyl-insoluble western blotting was used to investigate oxidative stress-induced pathological aggregation of TDP-43 in neuronal cells. N2a cells were transiently transfected with empty vector pcDNA3.1, pcDNA3.1-HA-PDI, or pcDNA3.1-HA-dnPDI according to the manufacturer’s instructions. After one-day culture and 1-h treatment with 1 mM H_2_O_2_ or PBS at 37 ℃, cells were harvested by centrifugation at 100 *g* at 4 °C for 5 min and ruptured on ice for 20 min with cell lysis buffer containing 1 × protease inhibitor cocktail (Target Mol). The cell lysates were then centrifuged at 17,000 *g* at 4 °C for 20 min to remove cell debris. The amount of loaded protein was normalized using a BCA Protein Quantification kit (Beyotime). A part of the supernatant was incubated with 1% sarkosyl for 30 min at room temperature. The mixture was then ultracentrifuged at 150,000 *g* for 30 min, and the supernatant was carefully removed and washed once with PBS. The sarkosyl-insoluble pellets were boiled in the SDS-PAGE loading buffer for 10 min. The remaining of the supernatant, which served as the total protein sample, was also boiled in the SDS-PAGE loading buffer for 10 min. The samples were separated by 12.5% SDS-PAGE and then western blotted. The sarkosyl-insoluble pellets from the above cells were probed with the anti-TDP-43 antibody, and the corresponding cell lysates were probed using the anti-TDP-43 antibody, anti-HA antibody, and anti-β-actin antibody, respectively, which served as the input controls. For calculating the amounts of sarkosyl-insoluble TDP-43, the ImageJ software (NIH) was used to assess the densitometry of TDP-43 bands, including insoluble full-length TDP-43 band and band of insoluble C-terminal TDP-43 fragment of 35 kDa. The normalized amount of insoluble TDP-43 aggregates in the above cells was determined as a ratio of the density of insoluble TDP-43 aggregate bands over that of the total TDP-43 bands in cell lysates. The following antibodies were used: rabbit anti-TDP-43 polyclonal antibody (Proteintech, 10782-2-AP, 1:2000), mouse anti-HA tag monoclonal antibody (Proteintech, 66006-2-Ig, 1:5000), mouse anti-β-actin monoclonal antibody (Proteintech, 66009-1-Ig, 1:5000), HRP-labeled goat anti-rabbit IgG (H+L) (Beyotime, A0208, 1:5000), and HRP-labeled goat anti-mouse IgG (H+L) (Beyotime, A0216, 1:5000).

### Immunogold electron microscopy of TDP-43 fibrils

N2a cells were transiently transfected with empty vector pcDNA3.1, pcDNA3.1-HA-PDI, or pcDNA3.1-HA-dnPDI according to the manufacturer’s instructions. After one-day culture and 1-h treatment with 1 mM H_2_O_2_ or PBS at 37 ℃, the above N2a cells were washed twice with cold PBS, digested with trypsin-EDTA solution (Beyotime, C0201), resuspended with 1 ml PBS, and centrifuged at 100 *g* for 5 min at 4 ℃ to harvest the cells. The cells were then resuspended with 450 μl of 10% sucrose solution in 10 mM Tris-HCl buffer (pH 7.4) containing 1 mM EGTA, 0.8 M NaCl, and 1 × protease inhibitor cocktail (Target Mol). The mixtures were sonicated 5 times on ice at 50 W and 5 s/5 s, and then centrifuged at 17,000 *g* for 20 min at 4 ℃ to remove the cell debris. A part of the supernatant was incubated with 1% sarkosyl for 30 min at room temperature. The mixture was then ultracentrifuged at 150,000 *g* for 30 min, and the supernatant was carefully removed and washed once with PBS. The sarkosyl-insoluble pellets were resuspended in PBS (50 μl). Sample aliquots of 10 μl were absorbed onto nickel grids for 1 min, and then washed three times with water for 30 s. Samples on grids were incubated with rabbit anti-TDP-43 polyclonal antibody (Proteintech, 10782-2-AP, 1:1000) for 30 min at room temperature and then blocked with 0.1% BSA in PBS for 15 min. 10-nm gold-labeled homologous secondary antibody (goat anti-rabbit IgG H&L, Abcam, ab27234, 1:20) was used to incubate the grids for 20 min at room temperature. Unbound gold-labeled homologous secondary antibodies were removed by washing with 200 μl of water drop by drop. Samples on grids were then stained with 2% (w/v) uranyl acetate for 1 min. The stained samples were examined using a JEM-1400 Plus transmission electron microscope (JEOL) operating at 100 kV.

### Annexin V-FITC apoptosis detection assay

Wild-type N2a cells and N2a cells stably expressing TDP-43 were transiently transfected with empty vector pcDNA3.1, pcDNA3.1-HA-PDI, or pcDNA3.1-HA-dnPDI. After one-day culture and with the treatment of 1 mM H_2_O_2_ or PBS at 37 ℃ for 1 h, apoptotic cells were detected by flow cytometry after staining with an annexin V-FITC apoptosis detection kit (4A Biotech, FXP018) according to the manufacturer’s instructions. In brief, the above N2a cells were harvested after digestion with trypsin-EDTA solution (Beyotime, C0201); the cells were washed twice with cold PBS at 100 *g* and 4 °C for 5 min and resuspended in 185 μl of binding buffer. Each sample was then incubated with 5 μl of annexin V-FITC and 10 μl of PI for 15 min at 4 °C in the dark and filtered through nylon sieves with screen mesh 300. Annexin V and PI binding were analyzed using an CytoFLEX flow cytometer (Beckman Coulter), and the percentage of apoptotic cells was calculated from the total number of cells (∼1 × 10^4^ cells) using CytExpert software. All apoptotic blot experiments were repeated at least three times.

### Ultrathin section TEM

N2a and HEK-293T cells stably expressing TDP-43 were transiently transfected with empty vector pcDNA3.1, pcDNA3.1-HA-PDI, or pcDNA3.1-HA-dnPDI. After one-day culture and 1-h treatment with 1 mM H_2_O_2_ or PBS at 37 ℃, cells were prefixed with 3% paraformaldehyde and 1.5% glutaraldehyde in PBS, then harvested and postfixed in 1% osmium tetroxide for 1 h using an ice bath; the samples were then dehydrated in graded acetone and embedded in 812 resins. Ultrathin sections of the cells were prepared using a Leica Ultracut S Microtome and negatively stained using 2% uranyl acetate and lead citrate. The doubly stained ultrathin sections of cells were examined using a JEM-1400 Plus transmission electron microscope (JEOL) operating at 100 kV. All experiments were further confirmed by biological repeats.

### Statistical Analysis

The data shown for each experiment were based on at least three technical replicates, as indicated in individual figure legends. Data are presented as mean ± S.D., and *P* values were determined using a two-sided Student’s *t*-test. Differences were considered statistically significant when *P* < 0.05. All experiments were further confirmed by biological repeats.

## Acknowledgments

We thank L. W. Ruddock (Faculty of Biochemistry and Molecular Medicine, University of Oulu), H.-N. Du (College of Life Sciences, Wuhan University), and Y. Liu (College of Life Sciences, Wuhan University) for the kind gifts of the human PDI plasmid, the human TDP-43 plasmid, and the G3BP1-DsRed plasmid, respectively; Z. Song and W. Zou (College of Life Sciences, Wuhan University) for their technical assistances with the TEM of ultrathin sections of cells; and L. Wang and C. C. Wang (Institute of Biophysics, Chinese Academy of Sciences) for their helpful suggestions.

## Author contributions

M.C. and Y.L. supervised the project. J.-Q.L., M.C. and Y.L. designed the experiments. J.-Q.L., H.L., L.-Q.W., K.W. and J.C. purified the human TDP-43 and the human PDI, and cultured the cells. J.-Q.L. and H.L. performed in vitro phase separation and droplet fusion experiments. J.-Q.L. performed western blotting, immunoblotting analyses, BiFC assay, immunofluorescence staining, immunogold EM, flow cytometry, and Ultrathin section TEM. X.L., Z.Y., X.X., Y.Z., W.L., and M.C. provided the brain and spinal cord samples. J.-Q.L. and Y. Li performed H&E staining of paraffin brain and spinal cord sections. Q.F. and J.Z. provided GSNO and SNO-PDI-producing methods. J.-Q.L., W.L., M.C. and Y.L. wrote the manuscript. All authors proofread and approved the manuscript.

## Data and materials availability

All data needed to evaluate the conclusions in the paper are present in the paper and/or the Supplementary Materials. Additional data related to this paper may be requested from the authors.

## Supplementary Materials

**table S1.**
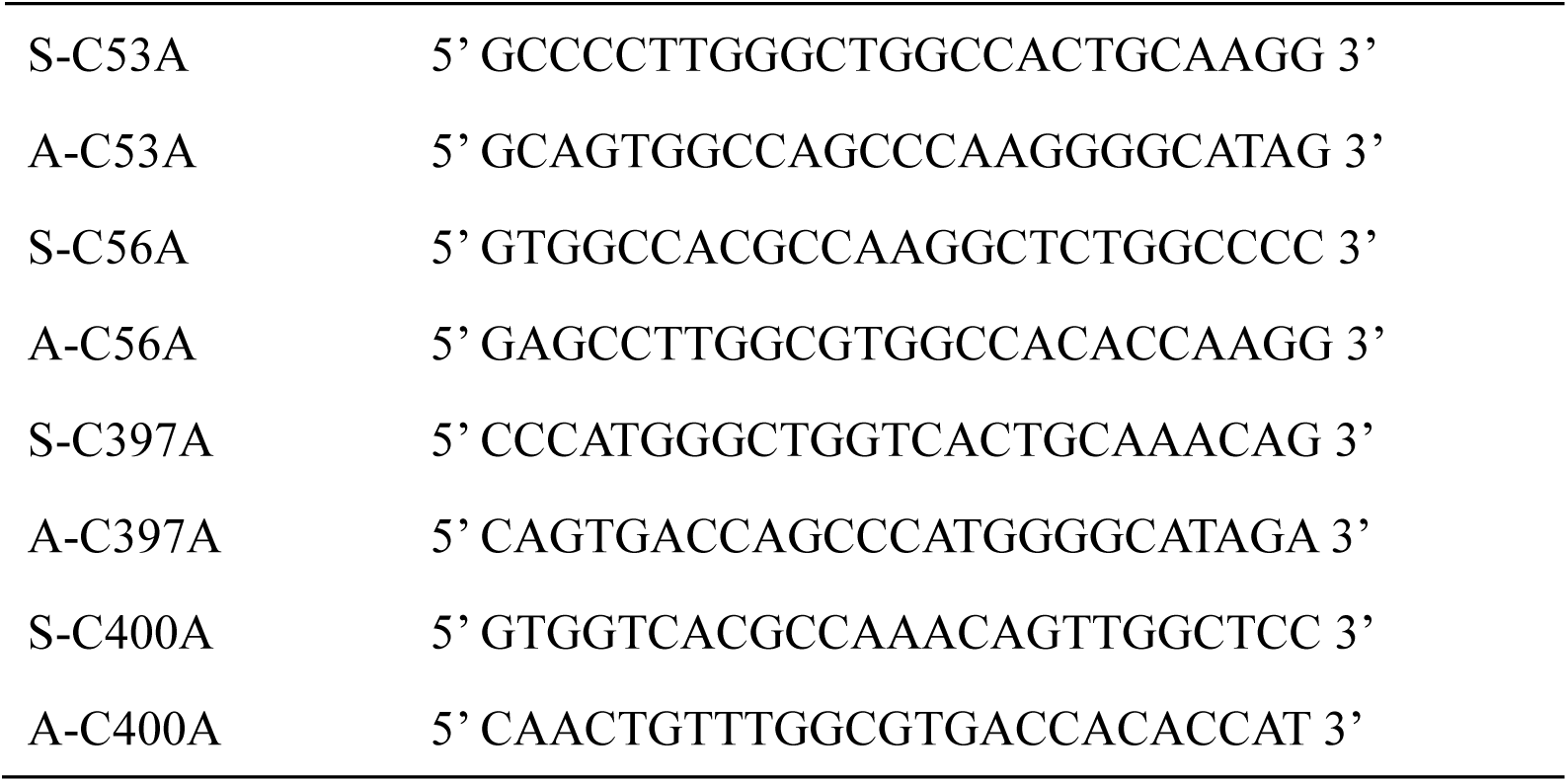
The primers designed for full-length human PDI with C53A/C56A/C397A/C400A mutation (dnPDI)

**figure S1.**
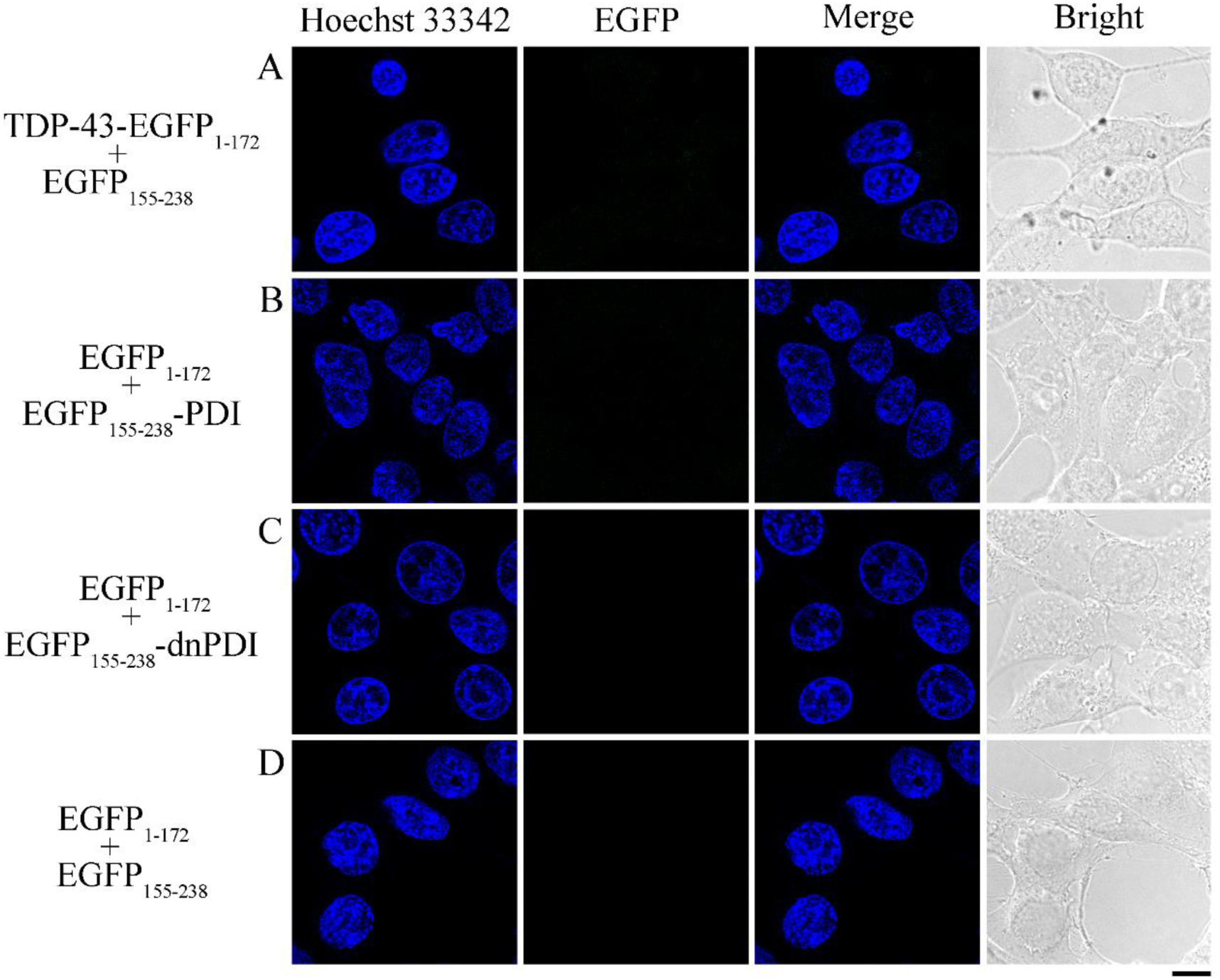
TDP-43 and PDI are required for this putative interaction in living cells. PDI includes wild-type PDI and its mutant dnPDI. (**A** to **D**) HEK-293T cells transiently expressing both full-length human TDP-43-EGFP_1-172_ and EGFP_155-238_ constructs (A) and HEK-293T cells transiently expressing both EGFP_1-172_ and EGFP_155-238_-wild-type PDI (B) or EGFP_155-238_-dnPDI (C) or EGFP_155-238_ (D) constructs were cultured for 1 day. Shown are nuclei stained with Hoechst 33342 (blue). EGFP (green) was not observed in (A to D). Scale bar, 10 μm.

**figure S2.**
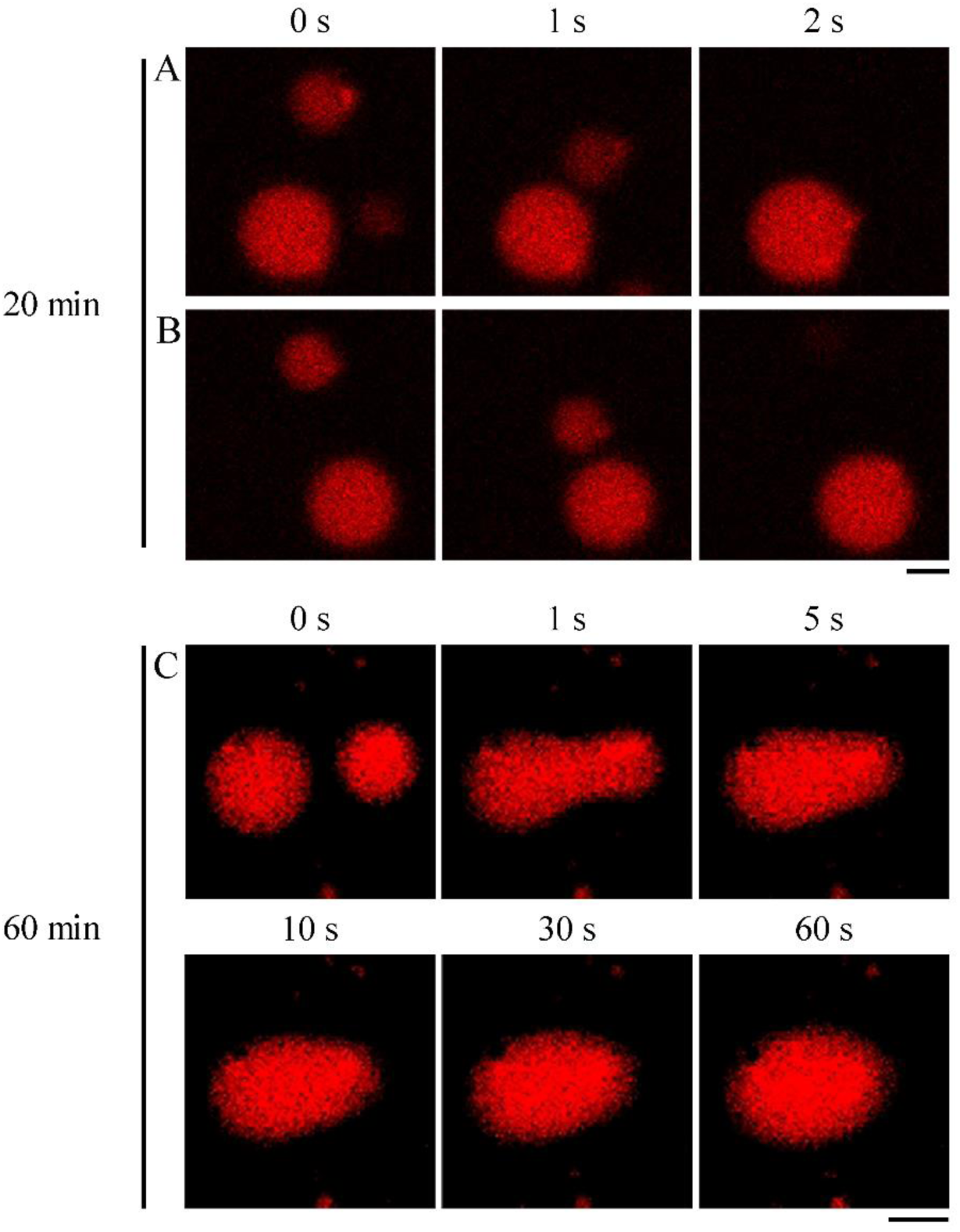
A droplet fusion event occurs in the regions of different time slots. (**A** and **B**) During the first 20 min after droplet initiation, two small liquid condensates of TDP-43 fused into one larger liquid droplet rapidly within 2 seconds. (**C**) After 60 min incubation, however, two small liquid condensates of TDP-43 fused into one larger liquid droplet slowly within 60 seconds. Fluorescence images of in vitro phase-separated droplets (red) of 10 μM TAMRA-labeled TDP-43 incubated with Tris-HCl buffer (pH 8.0) containing 10% PEG 3350 on ice for the indicated time. Scale bars, 2 μm.

**figure S3.**
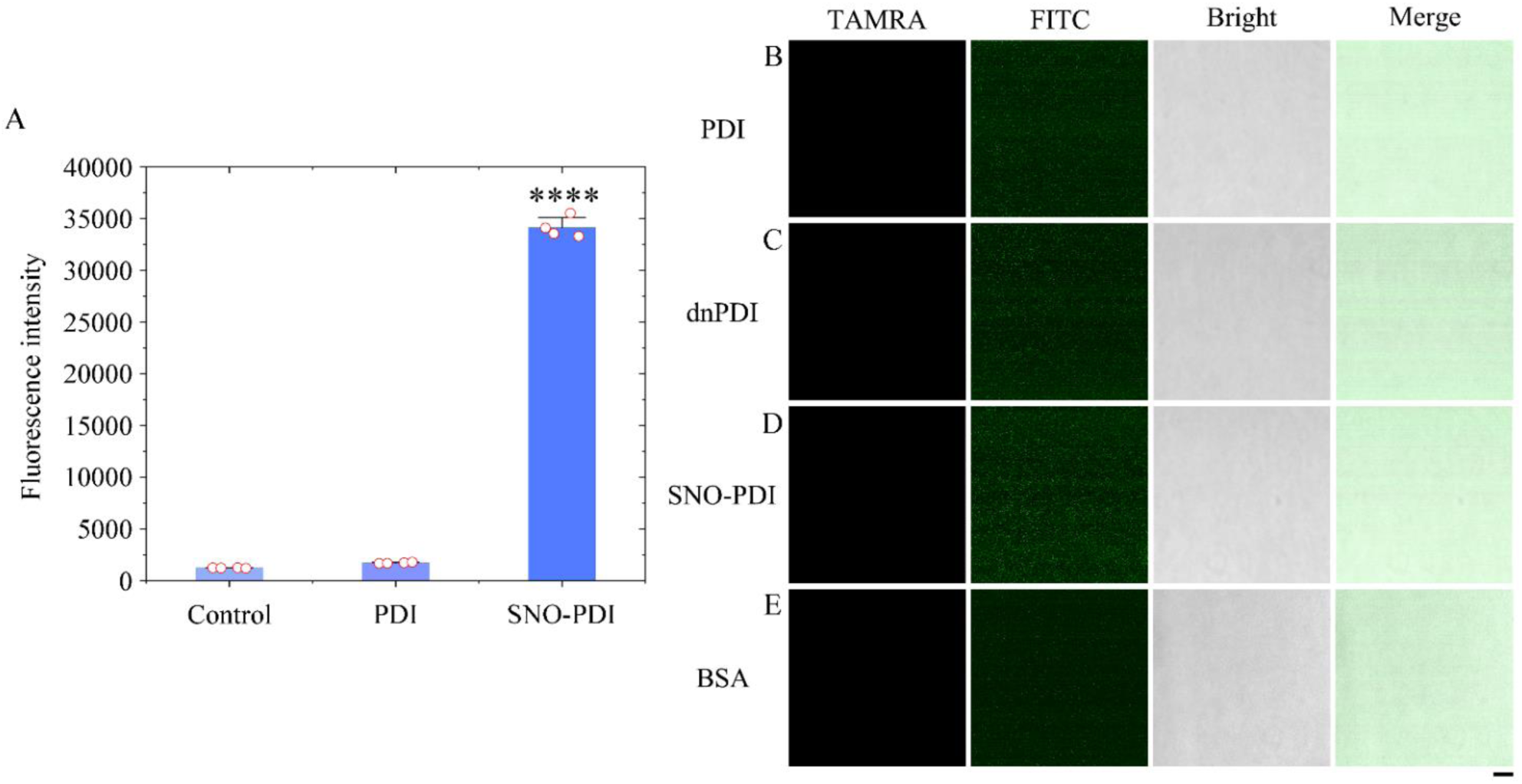
**(A) Characterization of S-nitrosylated PDI by fluorescence spectroscopy.** Tris-HCl buffer (pH 8.0) (control) and Tris-HCl buffer (pH 8.0) containing wild-type PDI or SNO-PDI were incubated with a solution containing 100 μM DAN and 100 μM CuSO_4_ in the dark at room temperature for 30 min. S-nitrosylated PDI was then assessed by the release of NO, causing the conversion of DAN into the fluorescent compound NAT. The fluorescence intensity of NAT in the aforementioned samples (open red circles shown in scatter plots) measured using excitation at 375 nm and emission at 450 nm, was expressed as the mean ± SD (with error bars) of values obtained in four independent experiments. SNO-PDI, *P* = 0.00000000088. Statistical analyses were performed using two-sided Student’s *t*-test. Values of *P* < 0.05 indicate statistically significant differences. The following notation is used throughout: \**P* < 0.05; \*\**P* < 0.01; \*\*\**P* < 0.001; and \*\*\*\**P* < 0.0001 relative to control (wild-type PDI). (**B** to **E) Wild-type PDI, dnPDI, SNO-PDI, or bovine serum albumin (BSA) alone did not form liquid droplets.** Fluorescence images of Tris buffer containing 10 μM FITC-labeled wild-type PDI (B), dnPDI (C), SNO-PDI (D), or BSA (E) (green) and 10% PEG 3350 on ice, as four controls for the regulation of TDP-43 LLPS by wild-type PDI. Scale bar, 10 μm.

**figure S4.**
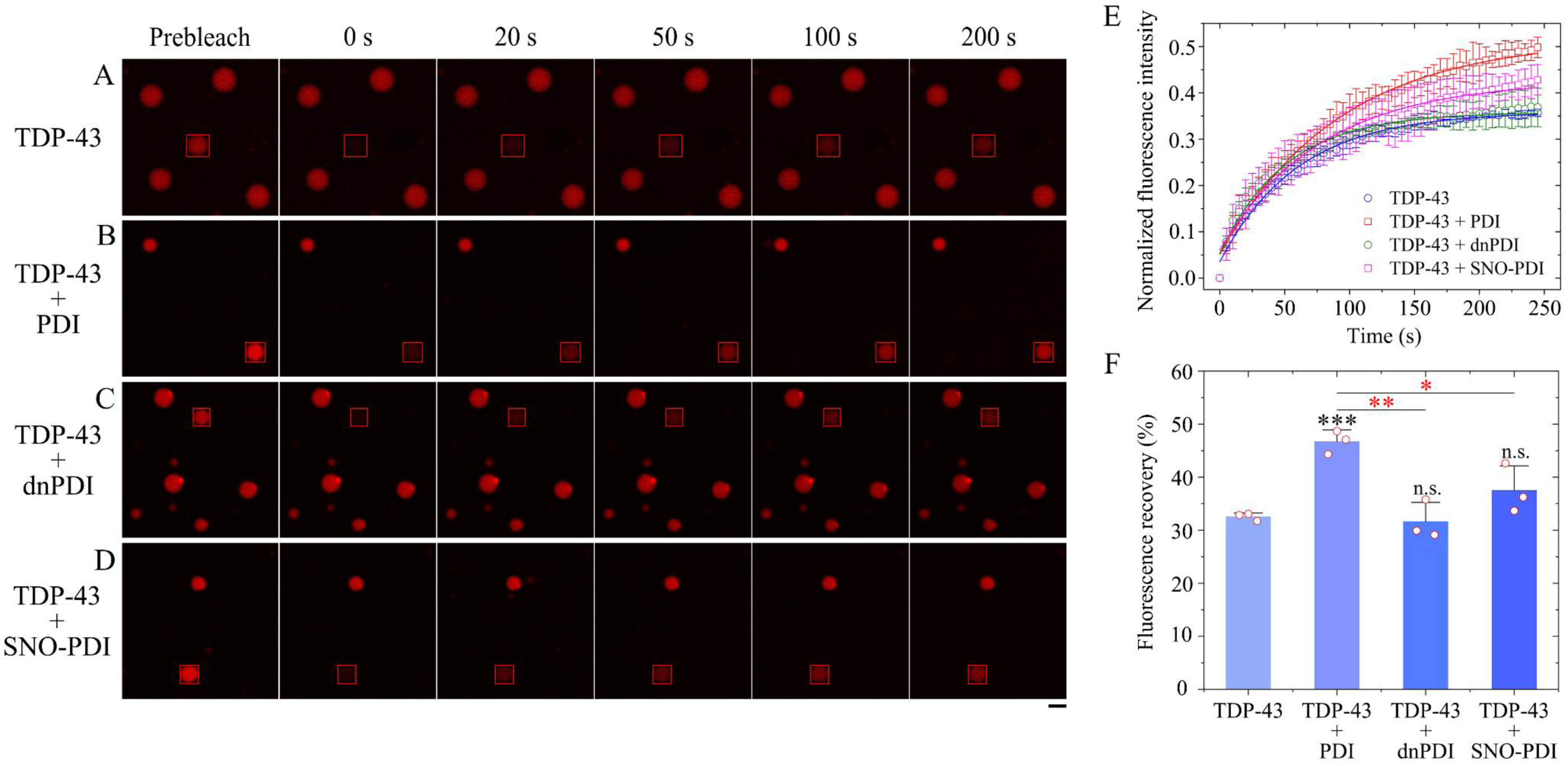
Wild-type PDI enhances fluorescence recovery and negatively modulates LLPS of TDP-43. (**A** to **D**) FRAP analysis on the selected liquid droplets of 10 μM TAMRA-labeled TDP-43 incubated with Tris-HCl buffer (pH 8.0) containing 10% PEG 3350 (A), 10 μM wild-type PDI and 10% PEG 3350 (B), 10 μM dnPDI and 10% PEG 3350 (C), or 10 μM SNO-PDI and 10% PEG 3350 (D) on ice before (prebleach), during (0 s), and after photobleaching (20, 50, 100, and 200 s, respectively). The internal photobleaching is marked by a red square. Scale bar, 5 μm. (**E**) Normalized kinetics of fluorescence recovery data of TDP-43 (blue circle), TDP-43 + wild-type PDI (red square), TDP-43 + dnPDI (olive circle), and TDP-43 + SNO-PDI (magenta square) obtained from FRAP intensity. The normalized fluorescence intensity is expressed as the mean ± SD (with error bars) of values obtained in three independent experiments. The solid lines show the best single exponential fit for the fluorescence intensity-time curves. (**F**) Fluorescence recovery percentage of TDP-43 condensates (open red circles shown in scatter plots) were expressed as the mean ± SD (with error bars) of values obtained in three biologically independent experiments. TDP-43 + wild-type PDI, *P* = 0.00045; TDP-43 + dnPDI, *P* = 0.681 or 0.00359; and TDP-43 + SNO-PDI, *P* = 0.139 or 0.0358. Statistical analyses were performed using two-sided Student’s *t*-test. Values of *P* < 0.05 indicate statistically significant differences. The following notation is used throughout: \**P* < 0.05; \*\**P* < 0.01; and \*\*\**P* < 0.001 relative to control (the fluorescence recovery for TDP-43 or TDP-43 incubated with wild-type PDI). n.s., no significance.

**figure S5.**
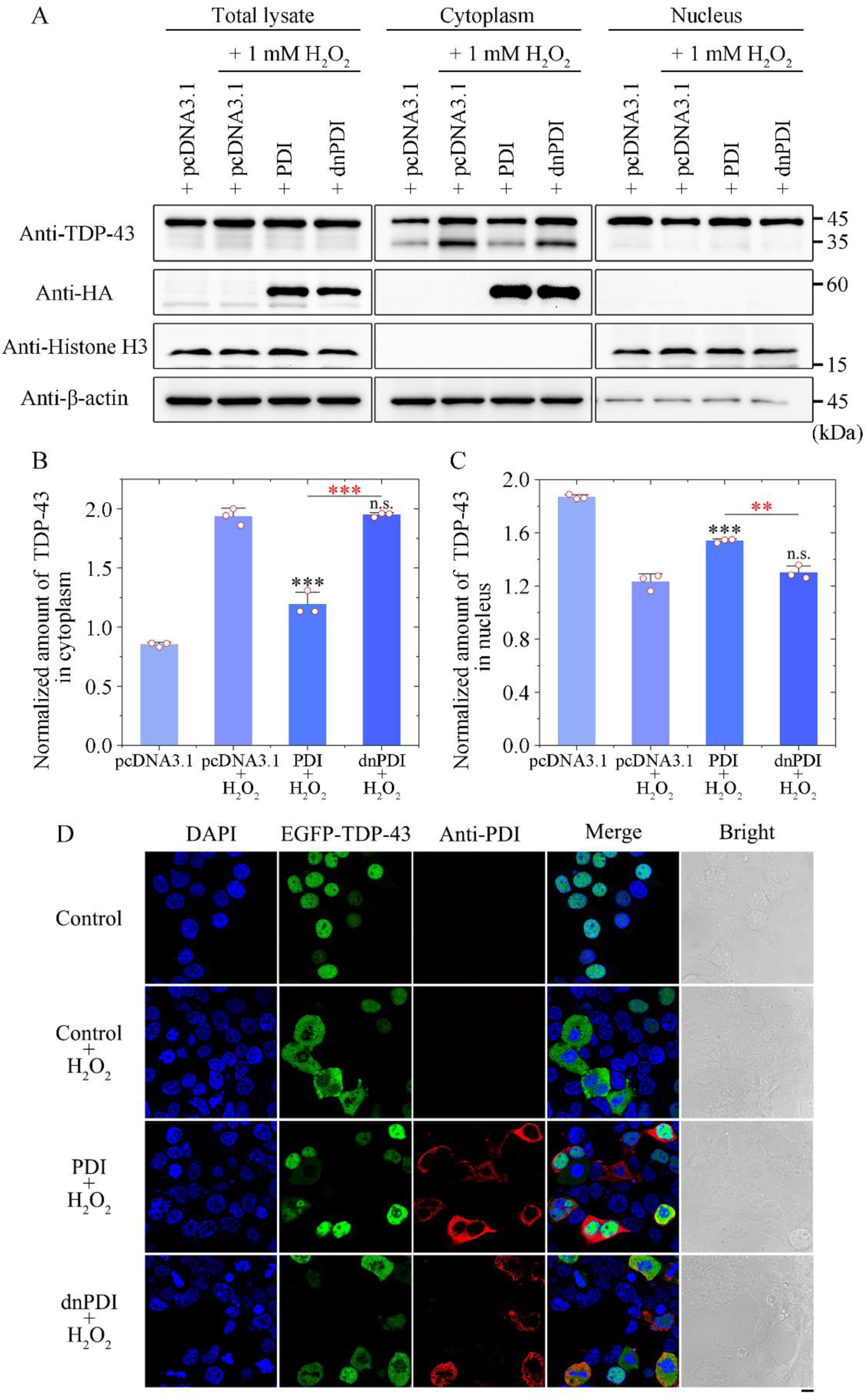
Wild-type PDI blocks the mislocalization of TDP-43 to the cytoplasm induced by H_2_O_2_. (**A**) Western blot for endogenous TDP-43 in the cytoplasmic and nuclear protein fractions and the corresponding cell lysates from HEK-293T cells transiently expressing HA-tagged wild-type PDI or HA-tagged dnPDI treated with 1 mM H_2_O_2_ for 1 h. β-actin and histone H3 served as markers for cytosol and nuclear fractions, respectively, and empty vector pcDNA3.1 as a control. (**B** and **C**) The normalized amount of endogenous TDP-43 in cytoplasm (B) and nucleus (C) of the aforementioned cells (open red circles shown in scatter plots) was expressed as the mean ± SD (with error bars) of values obtained in three independent experiments. Wild-type PDI + H_2_O_2_, *P* = 0.00049 (B) and 0.00099 (C); and dnPDI + H_2_O_2_, *P* = 0.741 or 0.00023 (B) and 0.200 or 0.0013 (C). Statistical analyses were performed using two-sided Student’s *t*-test. Values of *P* < 0.05 indicate statistically significant differences. The following notation is used throughout: \**P* < 0.05; \*\**P* < 0.01; and \*\*\**P* < 0.001 relative to control (pcDNA3.1 + H_2_O_2_ or wild-type PDI + H_2_O_2_). n.s., no significance. (**D**) Immunofluorescence imaging of HEK-293T cells transiently expressing both EGFP-TDP-43 and empty vector pcDNA3.1 (control), transiently expressing both EGFP-TDP-43 and pcDNA3.1 treated with 1 mM H_2_O_2_ for 1 h (control + H_2_O_2_), transiently expressing both EGFP-TDP-43 and wild-type PDI treated with 1 mM H_2_O_2_ for 1 h (wild-type PDI + H_2_O_2_), and transiently expressing both EGFP-TDP-43 and dnPDI also treated with 1 mM H_2_O_2_ for 1 h (dnPDI + H_2_O_2_), using antibody against PDI (red) and staining with DAPI (blue). EGFP-TDP-43 (green) was observed in (D). Scale bar, 10 μm.

**figure S6.**
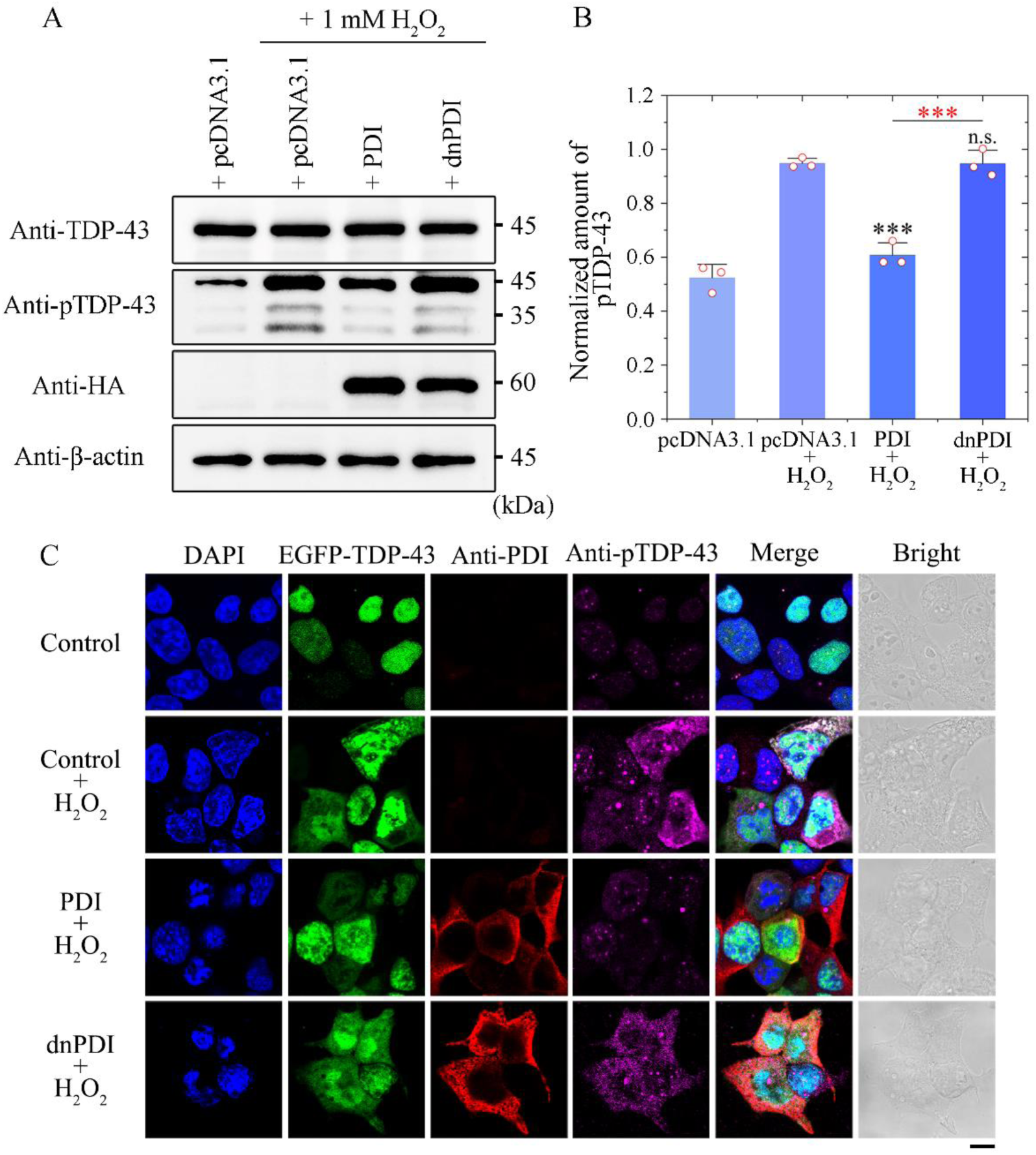
Wild-type PDI blocks pathological phosphorylation of TDP-43 induced by H_2_O_2_. (**A**) Western blot for phosphorylated endogenous TDP-43 (pTDP-43) in HEK-293T cells transiently expressing empty vector pcDNA3.1, transiently expressing pcDNA3.1 treated with 1 mM H_2_O_2_ for 1 h, transiently expressing wild-type PDI treated with 1 mM H_2_O_2_ for 1 h, and transiently expressing dnPDI also treated with 1 mM H_2_O_2_ for 1 h. The cell lysates from the aforementioned cells were probed by anti-TDP-43, anti-pTDP-43, anti-HA, and anti-β-actin antibodies. (**B**) The normalized amount of pTDP-43 in the aforementioned cells (open red circles shown in scatter plots) was expressed as the mean ± SD (with error bars) of values obtained in three independent experiments. Wild-type PDI + H_2_O_2_, *P* = 0.00027; and dnPDI + H_2_O_2_, *P* = 0.977 or 0.00095. Statistical analyses were performed using two-sided Student’s *t*-test. Values of *P* < 0.05 indicate statistically significant differences. The following notation is used throughout: \**P* < 0.05; \*\**P* < 0.01; and \*\*\**P* < 0.001 relative to control (pcDNA3.1 + H_2_O_2_ or wild-type PDI + H_2_O_2_). n.s., no significance. (**C**) Immunofluorescence imaging of HEK-293T cells transiently expressing both EGFP-TDP-43 and pcDNA3.1 (control), transiently expressing both EGFP-TDP-43 and pcDNA3.1 treated with 1 mM H_2_O_2_ for 1 h (control + H_2_O_2_), transiently expressing both EGFP-TDP-43 and wild-type PDI treated with 1 mM H_2_O_2_ for 1 h (wild-type PDI + H_2_O_2_), and transiently expressing both EGFP-TDP-43 and dnPDI also treated with 1 mM H_2_O_2_ for 1 h (dnPDI + H_2_O_2_), using antibodies against PDI (red) and pTDP-43 (magenta) and staining with DAPI (blue). EGFP-TDP-43 (green) was observed in (C). Scale bar, 10 μm.

**figure S7.**
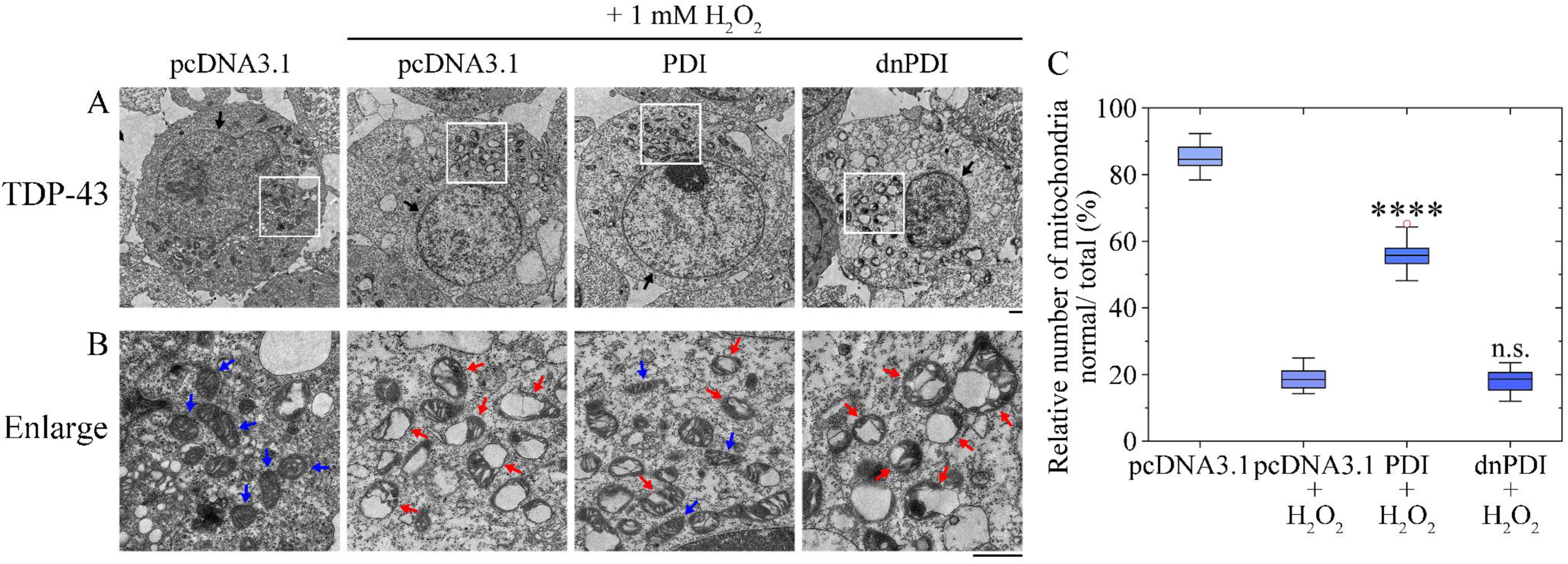
Wild-type PDI suppress mitochondrial damage resulting from H_2_O_2_-induced TDP-43 aggregation. (**A** and **B**) Wild-type PDI blocked H_2_O_2_-induced mitochondrial damage in HEK-293T cells stably expressing TDP-43. TEM imaging of stable HEK-293T cells transiently expressing empty vector pcDNA3.1, transiently expressing pcDNA3.1 treated with 1 mM H_2_O_2_ for 1 h, transiently expressing wild-type PDI treated with 1 mM H_2_O_2_ for 1 h, and transiently expressing dnPDI also treated with 1 mM H_2_O_2_ for 1 h (A). The enlarged regions (B) show 14-fold enlarged images from (A) and display the detailed structures of mitochondria in cells. Nuclei and normal mitochondria in HEK-293T cells are highlighted using black arrows and blue arrows, respectively. Abnormal mitochondria in cells with mitochondrial vacuolization are highlighted by red arrows. Samples were negatively stained using 2% uranyl acetate and lead citrate. Scale bars, 1 μm. (**C**) Wild-type PDI significantly decreased H_2_O_2_-induced mitochondrial damage in HEK-293T cells stably expressing TDP-43. Box plots analyzing the relative number of mitochondria (normal/total) in the aforementioned cells and showing the quantification of TEM images in *n* = 30 cells examined over 3 independent experiments. The experiments were done blind and average 19 to 28 mitochondria were present in each cell. The boxes (blue) extend from the 25th to 75th percentile (quantiles 1 and 3). Minima, maxima, centre, and bounds of box represent quantile 1 minus 1.5 × interquartile range, quantile 3 plus 1.5 × interquartile range, median (black line), and quantiles 1 and 3, respectively. Bounds of whiskers (minima and maxima) and outliers (open red circles). Wild-type PDI + H_2_O_2_, *P* = 1.1 × 10^-43^; and dnPDI + H_2_O_2_, *P* = 0.982. Statistical analyses were performed using two-sided Student’s *t*-test. Values of *P* < 0.05 indicate statistically significant differences. The following notation is used throughout: \**P* < 0.05; \*\**P* < 0.01; \*\*\**P* < 0.001; and \*\*\*\**P* < 0.0001 relative to control (pcDNA3.1 + H_2_O_2_). n.s., no significance.

## REFERENCES AND NOTES

1. T. J. Cohen, V. M. Y. Lee, J. Q. Trojanowski, TDP-43 functions and pathogenic mechanisms implicated in TDP-43 proteinopathies. Trends Mol. Med. 17, 659−667 (2011).

2. M. Neumann, D. Sampathu, L. Kwong, A. Truax, M. Micsenyi, T. Chou, J. Bruce, T. Schuck, M. Grossman, C. Clark, Ubiquitinated TDP-43 in frontotemporal lobar degeneration and amyotrophic lateral sclerosis. Science 314, 130−133 (2006).

3. X. Wang, Y. Hu, R. Xu, The pathogenic mechanism of TAR DNA-binding protein 43 (TDP-43) in amyotrophic lateral sclerosis. Neural Regen. Res. 19, 800−806 (2024).

4. Y. Iguchi, Y. Takahashi, J. Li, K. Araki, Y. Amakusa, Y. Kawakami, K. Kobayashi, S. Yokoi, M. Katsuno, IκB kinase phosphorylates cytoplasmic TDP-43 and promotes its proteasome degradation. J. Cell Biol. 223, e202302048 (2024).

5. A. E. Perlegos, J. Durkin, S. J. Belfer, A. Rodriguez, O. Shcherbakova, K. Park, J. Luong, N. M. Bonini, M. S. Kayser, TDP-43 impairs sleep in *Drosophila* through *Ataxin*-2-dependent metabolic disturbance. Sci. Adv. 10, eadj4457 (2024).

6. K. Sharma, F. Stockert, J. Shenoy, M. Berbon, M. B. Abdul-Shukkoor, B. Habenstein, A. Loquet, M. Schmidt, M. Fändrich, Cryo-EM observation of the amyloid key structure of polymorphic TDP-43 amyloid fibrils. Nat. Commun. 15, 486 (2024).

7. J. I. Ayers, D. R. Borchelt, Phenotypic diversity in ALS and the role of poly-conformational protein misfolding. Acta Neuropathol. 142, 41−55 (2021).

8. W. C. Xu, J. Z. Liang, C. Li, Z. X. He, H. Y. Yuan, B. Y. Huang, X. L. Liu, B. Tang, D. W. Pang, H. N. Du, Y. Yang, J. Chen, L. Wang, M. Zhang, Y. Liang, Pathological hydrogen peroxide triggers the fibrillization of wild-type SOD1 via sulfenic acid modification of Cys-111. Cell Death Dis. 9, 67 (2018).

9. L. Q. Wang, Y. Ma, H. Y. Yuan, K. Zhao, M. Y. Zhang, Q. Wang, X. Huang, W. C. Xu, B. Dai, J. Chen, D. Li, D. Zhang, Z. Wang, L. Zou, P. Yin, C. Liu, Y. Liang, Cryo-EM structure of an amyloid fibril formed by full-length human SOD1 reveals its conformational conversion. Nat. Commun. 13, 3491 (2022).

10. P. Foulds, E. McAuley, L. Gibbons, Y. Davidson, S. M. Pickering-Brown, D. Neary, J. S. Snowden, D. Allsop, D. M. A. Mann, TDP-43 protein in plasma may index TDP-43 brain pathology in Alzheimer’s disease and frontotemporal lobar degeneration. Acta Neuropathol. 116, 141−146 (2008).

11. W. T. Hu, K. A. Josephs, D. S. Knopman, B. F. Boeve, D. W. Dickson, R. C. Petersen, J. E. Parisi, Temporal lobar predominance of TDP-43 neuronal cytoplasmic inclusions in Alzheimer disease. Acta Neuropathol. 116, 215−220 (2008).

12. K. A. Josephs, M. E. Murray, J. L. Whitwell, J. E. Parisi, L. Petrucelli, C. R. Jack, R. C. Petersen, D. W. Dickson, Staging TDP-43 pathology in Alzheimer’s disease. Acta Neuropathol. 127, 441−450 (2014).

13. B. D. James, R. S. Wilson, P. A. Boyle, J. Q. Trojanowski, D. A. Bennett, J. A. Schneider, TDP-43 stage, mixed pathologies, and clinical Alzheimer’s-type dementia. Brain 139, 2983−2993 (2016).

14. Y. H. Shih, L. H. Tu, T. Y. Chang, K. Ganesan, W. W. Chang, P. S. Chang, Y. S. Fang, Y. T. Lin, L. W. Jin, Y. R. Chen, TDP-43 interacts with amyloid-β, inhibits fibrillization, and worsens pathology in a model of Alzheimer’s disease. Nat. Commun. 11, 5950 (2020).

15. A. Meneses, S. Koga, J. O’Leary, D. W. Dickson, G. Bu, N. Zhao, TDP-43 pathology in Alzheimer’s disease. Mol. Neurodegener. 16, 84 (2021).

16. A. F. Carlos, N. Tosakulwong, S. D. Weigand, M. L. Senjem, C. G. Schwarz, D. S. Knopman, B. F. Boeve, R. C. Petersen, A. T. Nguyen, R. R. Reichard, D. W. Dickson, C. R. Jack, V. Lowe, J. L. Whitwell, K. A. Josephs, TDP-43 pathology effect on volume and flortaucipir uptake in Alzheimer’s disease. Alzheimers Dement. 19, 2343−2354 (2023).

17. V. Estades Ayuso, S. Pickles, T. Todd, M. Yue, K. Jansen-West, Y. Song, J. González Bejarano, B. Rawlinson, M. DeTure, N. R. Graff-Radford, B. F. Boeve, D. S. Knopman, R. C. Petersen, D. W. Dickson, K. A. Josephs, L. Petrucelli, M. Prudencio, TDP-43-regulated cryptic RNAs accumulate in Alzheimer’s disease brains. Mol. Neurodegener. 18, 57 (2023).

18. A. R. Agra Almeida Quadros, Z. Li, X. Wang, I. S. Ndayambaje, S. Aryal, N. Ramesh, M. Nolan, R. Jayakumar, Y. Han, H. Stillman, C. Aguilar, H. J. Wheeler, T. Connors, J. Lopez-Erauskin, M. W. Baughn, Z. Melamed, M. S. Beccari, L. Olmedo Martinez, M. Canori, C. Z. Lee, L. Moran, I. Draper, A. S. Kopin, D. H. Oakley, D. W. Dickson, D. W. Cleveland, B. T. Hyman, S. Das, N. Ertekin-Taner, C. Lagier-Tourenne, Cryptic splicing of stathmin-2 and UNC13A mRNAs is a pathological hallmark of TDP-43-associated Alzheimer’s disease. Acta Neuropathol. 147, 9 (2024).

19. P. Kosuri, J. Alegre-Cebollada, J. Feng, A. Kaplan, A. Inglës-Prieto, C. L. Badilla, B. R. Stockwell, J. M. Sanchez-Ruiz, A. Holmgren, J. M. Fernández, Protein folding drives disulfide formation. Cell 151, 794−806 (2012).

20. C. Wang, W. Li, J. Ren, J. Fang, H. Ke, W. Gong, W. Feng, C. C. Wang, Structural insights into the redox-regulated dynamic conformations of human protein disulfide isomerase. Antioxid. Redox Signal. 19, 36−45 (2013).

21. T. Uehara, T. Nakamura, D. Yao, Z. Q. Shi, Z. Gu, Y. Ma, E. Masliah, Y. Nomura, S. A. Lipton, S-nitrosylated protein-disulphide isomerase links protein misfolding to neurodegeneration. Nature 441, 513−517 (2006).

22. A. K. Walker, M. A. Farg, C. R. Bye, C. A. McLean, M. K. Horne, J. D. Atkin, Protein disulphide isomerase protects against protein aggregation and is S-nitrosylated in amyotrophic lateral sclerosis. Brain 133, 105−116 (2010).

23. S. Parakh, S. Shadfar, E. R. Perri, A. M. G. Ragagnin, C. V. Piattoni, M. B. Fogolín, K. C. Yuan, H. Shahheydari, E. K. Don, C. J. Thomas, Y. Hong, M. A. Comini, A. S. Laird, D. M. Spencer, J. D. Atkin, The redox activity of protein disulfide isomerase inhibits ALS phenotypes in cellular and Zebrafish models. iScience 23, 101097 (2020).

24. D. B. Medinas, P. Rozas, C. Hetz, Critical roles of protein disulfide isomerases in balancing proteostasis in the nervous system. J. Biol. Chem. 298, 102087 (2022).

25. Y. Tabata, K. Takano, T. Ito, M. Iinuma, T. Yoshimoto, H. Miura, Y. Kitao, S. Ogawa, O. Hori, Vaticanol B, a resveratrol tetramer, regulates endoplasmic reticulum stress and inflammation. Am. J. Physiol. Cell Physiol. 293, C411–C418 (2007).

26. C. Turano, S. Coppari, F. Altieri, A. Ferraro, Proteins of the PDI family: Unpredicted non-ER locations and functions. J. Cell. Physiol. 193, 154−163 (2002).

27. A. Igbaria, P. I. Merksamer, A. Trusina, F. Tilahun, J. R. Johnson, O. Brandman, N. J. Krogan, J. S. Weissman, F. R. Papa, Chaperone-mediated reflux of secretory proteins to the cytosol during endoplasmic reticulum stress, Proc. Natl. Acad. Sci. U.S.A 116, 11291−11298 (2019).

28. D. Sicari, F. G. Centonze, R. Pineau, P. J. Le Reste, L. Negroni, S. Chat, M. A. Mohtar, D. Thomas, R. Gillet, T. Hupp, E. Chevet, A. Igbaria, Reflux of endoplasmic reticulum proteins to the cytosol inactivates tumor suppressors. EMBO Rep. 22, e51412 (2021).

29. A. D. da Purificação, V. Debbas, L. Y. Tanaka, G. V. de Mello Gabriel, J. Wosniak Júnior, T. C. De Bessa, S. Garcia-Rosa, F. R. Martins Laurindo, P. V. Santos Oliveira, DNAJB12 and DNJB14 are non-redundant Hsp40 redox chaperones involved in endoplasmic reticulum protein reflux. Biochim. Biophys. Acta Gen. Subj. 1868, 130502 (2024).

30. L. P. Bergeron-Sandoval, N. Safae., S. W. Michnick, Mechanisms and consequences of macromolecular phase separation. Cell 165, 1067−1079 (2016).

31. S. Boeynaems, S. Alberti, N. L. Fawzi, T. Mittag, M. Polymenidou, F. Rousseau, J. Schymkowitz, J. Shorter, B. Wolozin, L. Van Den Bosch, P. Tompa, M. Fuxreiter, Protein phase separation: a new phase in cell biology. Trends Cell Biol. 28, 420−435 (2018).

32. S. F. Shimobayashi, P. Ronceray, D. W. Sanders, M. P. Haataja, C. P. Brangwynne, Nucleation landscape of biomolecular condensates. Nature 599, 503−506 (2021).

33. Y. Shin, C. P. Brangwynne, Liquid phase condensation in cell physiology and disease. Science 357, eaaf4382 (2017).

34. S. Maharana, J. Wang, D. K. Papadopoulos, D. Richter, A. Pozniakovsky, I. Poser, M. Bickle, S. Rizk, J. Guillén-Boixet, T. M. Franzmann, M. Jahnel, L. Marrone, Y. T. Chang, J. Sterneckert, P. Tomancak, A. A. Hyman, S. Alberti, RNA buffers the phase separation behavior of prion-like RNA binding proteins. Science 360, 918−921 (2018).

35. F. Gasset-Rosa, S. Lu, H. Yu, C. Chen, Z. Melamed, L. Guo, J. Shorter, S. Da Cruz, D. W. Cleveland, Cytoplasmic TDP-43 de-mixing independent of stress granules drives inhibition of nuclear import, loss of nuclear TDP-43, and cell death. Neuron 102, 339−357 (2019).

36. W. M. Babinchak, B. K. Dumm, S. Venus, S. Boyko, A. A. Putnam, E. Jankowsky, W. K. Surewicz, Small molecules as potent biphasic modulators of protein liquid−liquid phase separation. Nat. Commun. 11, 5574 (2020).

37. H. Yu, S. Lu, K. Gasior, D. Singh, S. Vazquez-Sanchez, O. Tapia, D. Toprani, M. S. Beccari, J. R. Yates III, S. Da Cruz, J. M. Newby, M. Lafarga, A. S. Gladfelter, E. Villa, D. W. Cleveland, HSP70 chaperones RNA-free TDP-43 into anisotropic intranuclear liquid spherical shells. Science 371, eabb4309 (2021).

38. L. A. Gruijs da Silva, F. Simonetti, S. Hutten, H. Riemenschneider, E. L. Sternburg, L. M. Pietrek, J. Gebel, V. Dötsch, D. Edbauer, G. Hummer, L. S. Stelzl, D. Dormann, Disease-linked TDP-43 hyperphosphorylation suppresses TDP-43 condensation and aggregation. EMBO J. 41, e108443 (2022).

39. S. Lu, J. Hu, O. A. Arogundade, A. Goginashvili, S. Vazquez-Sanchez, J. K. Diedrich, J. Gu, J. Blum, S. Oung, Q. Ye, H. Yu, J. Ravits, C. Liu, J. R. Yates III, D. W. Cleveland, Heat-shock chaperone HSPB1 regulates cytoplasmic TDP-43 phase separation and liquid-to-gel transition. Nat. Cell Biol. 24, 1378−1393 (2022).

40. J. Tao, Y. Zeng, B. Dai, Y. Liu, X. Pan, L. Q. Wang, J. Chen, Y. Zhou, Z. Lu, L. Xie, Y. Liang, Excess PrP^C^ inhibits muscle cell differentiation via miRNA-enhanced liquid−liquid phase separation implicated in myopathy. Nat. Commun. 14, 8131 (2023).

41. X. Zhou, L. Sumrow, K. Tashiro, L. Sutherland, D. Liu, T. Qin, M. Kato, G. Liszczak, S. L. McKnight, Mutations linked to neurological disease enhance self-association of low-complexity protein sequences. Science 377, eabn5582 (2022).

42. J. R. Mann, A. M. Gleixner, J. C. Mauna, E. Gomes, M. R. DeChellis-Marks, P. G. Needham, K. E. Copley, B. Hurtle, B. Portz, N. J. Pyles, L. Guo, C. B. Calder, Z. P. Wills, U. B. Pandey, J. K. Kofler, J. L. Brodsky, A. Thathiah, J. Shorter, C. J. Donnelly, RNA binding antagonizes neurotoxic phase transitions of TDP-43. Neuron 102, 321−338 (2019).

43. J. G. Morato, F. Hans, F. von Zweydorf, R. Feederle, S. J. Elsässer, A. A. Skodras, C. J. Gloeckner, E. Buratti, M. Neumann, P. J. Kahle, Sirtuin-1 sensitive lysine-136 acetylation drives phase separation and pathological aggregation of TDP-43. Nat. Commun. 13, 1223 (2022).

44. A. E. Conicella, G. L. Dignon, G. H. Zerze, H. B. Schmidt, A. M. D’Ordine, Y. C. Kim, R. Rohatgi, Y. M. Ayala, J. Mittal, N. L. Fawzi, TDP-43 α-helical structure tunes liquid–liquid phase separation and function. Proc. Natl. Acad. Sci. U.S.A 117, 5883−5894 (2020).

45. A. Wang, A. E. Conicella, H. B. Schmidt, E. W. Martin, S. N. Rhoads, A. N. Reeb, A. Nourse, D. R. Montero, V. H. Ryan, R. Rohatgi, F. Shewmaker, M. T. Naik, T. Mittag, Y. M. Ayala, N. L. Fawzi, A single N-terminal phosphomimic disrupts TDP-43 polymerization, phase separation, and RNA splicing. EMBO J. 37, e97452 (2018).

46. X. Zuo, J. Zhou, Y. Li, K. Wu, Z. Chen, Z. Luo, X. Zhang, Y. Liang, M. A. Esteban, Y. Zhou, X. D. Fu, TDP-43 aggregation induced by oxidative stress causes global mitochondrial imbalance in ALS. Nat. Struct. Mol. Biol. 28, 132−142 (2021).

47. Y. Iguchi, M. Katsuno, S. Takagi, S. Ishigaki, J. i. Niwa, M. Hasegawa, F. Tanaka, G. Sobue, Oxidative stress induced by glutathione depletion reproduces pathological modifications of TDP-43 linked to TDP-43 proteinopathies. Neurobiol. Dis. 45, 862–870 (2012).

48. L. Kang, M. Piao, N. Liu, W. Gu, C. Feng, Sevoflurane exposure induces neuronal cell ferroptosis initiated by increase of intracellular hydrogen peroxide in the developing brain via ER stress ATF3 activation. Mol. Neurobiol. 61, in press (2024). doi: 10.1007/s12035-023-03695-z

49. C. D. Hu, Y. Chinenov, T. K. Kerppola, Visualization of interactions among bZIP and Rel family proteins in living cells using bimolecular fluorescence complementation. Mol. Cell 9, 789−798 (2002).

50. C. D. Hu, T. K. Kerppola, Simultaneous visualization of multiple protein interactions in living cells using multicolor fluorescence complementation analysis. Nat. Biotechnol. 21, 539−545 (2003).

51. S. Alberti, A. Gladfelter, T. Mittag, Considerations and challenges in studying liquid-liquid phase separation and biomolecular condensates. Cell 176, 419−434 (2019).

52. B. Dai, T. Zhong, Z. X. Chen, W. Chen, N. Zhang, X. L. Liu, L. Q. Wang, J. Chen, Y. Liang, Myricetin slows liquid-liquid phase separation of Tau and activates ATG5-dependent autophagy to suppress Tau toxicity. J. Biol. Chem. 297, 101222 (2021).

53. Y. Y. Gao, T. Zhong, L. Q. Wang, N. Zhang, Y. Zeng, J. Y. Hu, H. B. Dang, J. Chen, Y. Liang, Zinc enhances liquid-liquid phase separation of Tau protein and aggravates mitochondrial damages in cells. Int. J. Biol. Macromol. 209, 703−715 (2022).

54. X. N. Li, Y. Gao, Y. Li, J. X. Yin, C. W. Yi, H. Y. Yuan, J. J. Huang, L. Q. Wang, J. Chen, Y. Liang, Arg177 and Asp159 from dog prion protein slow liquid–liquid phase separation and inhibit amyloid formation of human prion protein. J. Biol. Chem. 299, 105329 (2023).

55. S. C. Barber, P. J. Shaw, Oxidative stress in ALS: key role in motor neuron injury and therapeutic target. Free Radical Biol. Med. 48, 629−641 (2010).

56. E. V. Ilieva, V. Ayala, M. Jové, E. Dalfó, D. Cacabelos, M. Povedano, M. J. Bellmunt, I. Ferrer, R. Pamplona, M. Portero-Otín, Oxidative and endoplasmic reticulum stress interplay in sporadic amyotrophic lateral sclerosis. Brain 130, 3111−3123 (2007).

57. P. Yang, C. Mathieu, R. M. Kolaitis, P. Zhang, J. Messing, U. Yurtsever, Z. Yang, J. Wu, Y. Li, Q. Pan, J. Yu, E. W. Martin, T. Mittag, H. J. Kim, J. P. Taylor, G3BP1 is a tunable switch that triggers phase separation to assemble stress granules. Cell 181, 325−345 (2020).

58. Y. Gwon, B. A. Maxwell, R. M. Kolaitis, P. Zhang, H. J. Kim, J. P. Taylor, Ubiquitination of G3BP1 mediates stress granule disassembly in a context-specific manner. Science 372, eabf6548 (2021).

59. C. Yang, Z. Wang, Y. Kang, Q. Yi, T. Wang, Y. Bai, Y. Liu, Stress granule homeostasis is modulated by TRIM21-mediated ubiquitination of G3BP1 and autophagy-dependent elimination of stress granules. Autophagy 19, 1934−1951 (2023).

60. B. D. Freibaum, J. Messing, H. Nakamura, U. Yurtsever, J. Wu, H. J. Kim, J. Hixon, R. M. Lemieux, J. Duffner, W. Huynh, K. Wong, M. White, C. Lee, R. E. Meyers, R. Parker, J. P. Taylor, Identification of small molecule inhibitors of G3BP-driven stress granule formation. J. Cell Biol. 223, e202308083 (2024).

61. M. Polymenidou, C. Lagier-Tourenne, K. R. Hutt, S. C. Huelga, J. Moran, T. Y. Liang, S. C. Ling, E. Sun, E. Wancewicz, C. Mazur, H. Kordasiewicz, Y. Sedaghat, J. P. Donohue, L. Shiue, C. F. Bennett, G. W. Yeo, D. W. Cleveland, Long pre-mRNA depletion and RNA missplicing contribute to neuronal vulnerability from loss of TDP-43. Nat. Neurosci. 14, 459−468 (2011).

62. M. Jo, S. Lee, Y. M. Jeon, S. Kim, Y. Kwon, H. J. Kim, The role of TDP-43 propagation in neurodegenerative diseases: integrating insights from clinical and experimental studies. Exp. Mol. Med. 52, 1652−1662 (2020).

63. B. A. Keller, K. Volkening, C. A. Droppelmann, L. C. Ang, R. Rademakers, M. J. Strong, Co-aggregation of RNA binding proteins in ALS spinal motor neurons: evidence of a common pathogenic mechanism. Acta Neuropathol. 124, 733−747 (2012).

64. L. T. W. Lin, A. Razzaq, S. E. Di Gregorio, S. Hong, B. Charles, M. H. Lopes, F. Beraldo, V. F. Prado, M. A. M. Prado, M. L. Duennwald, Hsp90 and its co-chaperone Sti1 control TDP-43 misfolding and toxicity. FASEB J. 35, e21594 (2021).

65. K. Luo, Z. Wang, K. Zhuang, S. Yuan, F. Liu, A. Liu, Suberoylanilide hydroxamic acid suppresses axonal damage and neurological dysfunction after subarachnoid hemorrhage via the HDAC1/HSP70/TDP-43 axis. Exp. Mol. Med. 54, 1423−1433 (2022).

66. E. Feneberg, D. Gordon, A. G. Thompson, M. J. Finelli, R. Dafinca, A. Candalija, P. D. Charles, I. Mäger, M. J. Wood, R. Fischer, B. M. Kessler, E. Gray, M. R. Turner, K. Talbot, An ALS-linked mutation in TDP-43 disrupts normal protein interactions in the motor neuron response to oxidative stress. Neurobiol. Dis. 144, 105050 (2020).

67. Y. Honjo, S. Kaneko, H. Ito, T. Horibe, M. Nagashima, M. Nakamura, K. Fujita, R. Takahashi, H. Kusaka, K. Kawakami, Protein disulfide isomerase-immunopositive inclusions in patients with amyotrophic lateral sclerosis. Amyotroph. Lateral Scler. 12, 444−450 (2011).

68. M. Vendruscolo, M. Fuxreiter, Protein condensation diseases: therapeutic opportunities. Nat. Commun. 13, 5550 (2022).

69. A. Patil, A. R. Strom, J. A. Paulo, C. K. Collings, K. M. Ruff, M. K. Shinn, A. Sankar, K. S. Cervantes, T. Wauer, J. D. St. Laurent, G. Xu, L. A. Becker, S. P. Gygi, R. V. Pappu, C. P. Brangwynne, C. Kadoch, A disordered region controls cBAF activity via condensation and partner recruitment. Cell 186, 4936−4955 (2023).

70. M. V. Vega, A. Nigro, S. Luti, C. Capitini, G. Fani, L. Gonnelli, F. Boscaro, F. Chiti, Isolation and characterization of soluble human full-length TDP-43 associated with neurodegeneration. FASEB J. 33, 10780–10793 (2019).

71. K. Wang, J. Q. Liu, T. Zhong, X. L. Liu, Y. Zeng, X. Qiao, T. Xie, Y. Chen, Y. Y. Gao, B. Tang, J. Li, J. Zhou, D. W. Pang, J. Chen, C. Chen, Y. Liang, Phase separation and cytotoxicity of Tau are modulated by protein disulfide isomerase and S-nitrosylation of this molecular chaperone. J. Mol. Biol. 432, 2141−2163 (2020).

72. W. M. Jones, J. B. Tapia, R. R. Tuttle, M. M. Reynolds, Thermogravimetric analysis and mass spectrometry allow for determination of chemisorbed reaction products on metal organic frameworks. Langmuir 36, 3903−3911 (2020).

73. Z. Gu, M. Kaul, B. Yan, S. J. Kridel, J. Cui, A. Strongin, J. W. Smith, R. C. Liddington, S. A. Lipton, S-Nitrosylation of matrix metalloproteinases: signaling pathway to neuronal cell death. Science 297, 1186−1190 (2002).

74. J. Haendeler, J. Hoffmann, V. Tischler, B. C. Berk, A. M. Zeiher, S. Dimmeler, Redox regulatory and anti-apoptotic functions of thioredoxin depend on S-nitrosylation at cysteine 69. Nat. Cell Biol. 4, 743−749 (2002).

